# DNA end sensing and cleavage by the Shedu anti-phage defense system

**DOI:** 10.1101/2023.08.10.552762

**Authors:** Luuk Loeff, Alexander Walter, Gian Tizio Rosalen, Martin Jinek

## Abstract

The detection of molecular patterns associated with invading pathogens is a hallmark of innate immune systems. Prokaryotes deploy sophisticated host-defense mechanisms in innate anti-phage immunity. Shedu is a single-component defense system comprising a putative nuclease SduA. Here we report cryo-EM structures of apo- and dsDNA-bound tetrameric SduA assemblies, revealing that the N-terminal domains of SduA form a clamp that recognizes free DNA ends. End binding positions the DNA over the PD/ExK nuclease domain, resulting in dsDNA nicking at a fixed distance from the 5’ end. The end-directed DNA nicking activity of Shedu prevents propagation of linear DNA *in vivo*. Finally, we show that phages escape Shedu immunity by suppressing their recombination-dependent DNA replication pathway. Taken together, these results define the antiviral mechanism of Shedu systems, underlining the paradigm that recognition of pathogen-specific nucleic acid structures is a conserved feature of innate immunity across all domains of life.

## Introduction

The evolutionary arms race between prokaryotes and invading mobile genetic elements has resulted in the emergence of a myriad of anti-viral defense systems that cluster in defense islands within host genomes^1^. This intrinsic feature of prokaryotic immune systems has facilitated systematic discovery of novel defense systems by identifying clusters of unknown genes adjacent to known defense operons^2–4^. Using this approach, numerous putative defense systems have recently been identified, including BREX^5^, DISARM^6^, Septu^2^, RADAR^3^, and Mokosh^4^, whose protein components are associated with diverse enzymatic activities. These “innate” immune systems are thought to provide multi-layered host defense, complementing the activities of canonical defense mechanisms such as restriction-modification, abortive infection, and adaptive immune systems such as CRISPR-Cas^7,8^.

For a small subset of these innate systems, the molecular triggers and mechanisms that underpin immunity have been uncovered^9–16^. For example, the CBASS system provides immunity by detecting a highly structured phage RNA^17^, resulting in the production of cyclic dinucleotides^18,19^ that subsequently activate downstream effector proteins to trigger the death of the infected host cell^18,20^. In contrast to CBASS, AVAST and CapRel^SJ46^ activate their downstream effectors by recognizing highly conserved phage proteins, such as portal, terminase, and the major capsid protein, respectively, to abort phage infection^21,22^. Although the immunological roles for most other novel defense systems have been determined, the molecular features recognized by these systems remain elusive, as do the functional mechanisms that underpin immunity.

The prokaryotic defense system Shedu encompasses a single gene *sduA* whose protein product is characterized by the presence of a hallmark domain (DUF4263) with predicted nuclease activity^2^. Homology searches reveal similarity to the RES domain of type III restriction-modification enzymes, which have been associated with nuclease and ATPase activities^23,24^. SduA displays distant homology to NucS/EndoMS, an archaeal RecB-like single stranded DNA (ssDNA) endonuclease that is involved in DNA mismatch repair^25,26^, pointing towards a functional mechanism in which Shedu directly targets the nucleic acids of invasive genetic elements.

In this study, we employed an integrative approach that combines single-particle cryo-electron microscopy (cryo-EM), biochemistry and phage infection assays to elucidate the molecular mechanism that underlies anti-phage defense by Shedu. Our biochemical and structural analysis of the Shedu system from *Escherichia coli* shows that SduA assembles into homotetrameric complex through its hallmark DUF4263 domain, which exhibits nickase activity on double stranded DNA substates (dsDNA). A cryo-EM structure of DNA-bound SduA, along with complementary biochemical assays reveal that Shedu systems recognize free ends of linear DNA molecules, poising the DNA for nicking at a fixed distance from the end. These results reveal that linear DNA ends are the molecular pattern recognized by Shedu systems, enabling phage sensing and nickase-based interference while ensuring self-vs-non-self discrimination.

## Results

### Shedu systems have distinct architectures

To understand the diversity of Shedu systems, we performed a phylogenetic analysis of 645 SduA orthologs, which grouped into two clusters that have distinct domain architectures based on their size and position of the signature domain DUF4263 (**Figure 1A**). For the first cluster, hereafter called type I, the DUF4263 is positioned in the C-terminal part of the protein coding sequence and preceded by a long N-terminal region (**Figure 1A** and **S1A**). By contrast, the second cluster, hereafter called type II, comprises orthologs with short extensions at either the N- or C-termini of the DUF4263 (**Figure 1A** and **S1A-B**). A fraction of 79% of Shedu systems belong to type I (**Figure 1B**), and have a mean size of 386 amino acids (**Figure 1C**), while type II systems account for the remaining 21% of systems (**Figure 1B**), and encode for miniature SduA proteins with a mean size of 254 amino acids (**Figure 1C**). Given the presence of subclusters in the type I clade (**Figure 1A**), we further performed sequence alignments to better understand its diversity. A sequence alignment of all type I systems displayed high sequence conservation of DUF4263, whereas the N-terminal regions showed low sequence conservation (**Figure S1C**). This indicates that the N-terminal regions of type I SduA proteins are highly diverse, suggesting that type I Shedu systems may provide distinct modes of defense. Taken together, this analysis suggests that Shedu constitute highly diverse immune systems that may elicit immunity through a variety of mechanisms.

**Figure 1:**
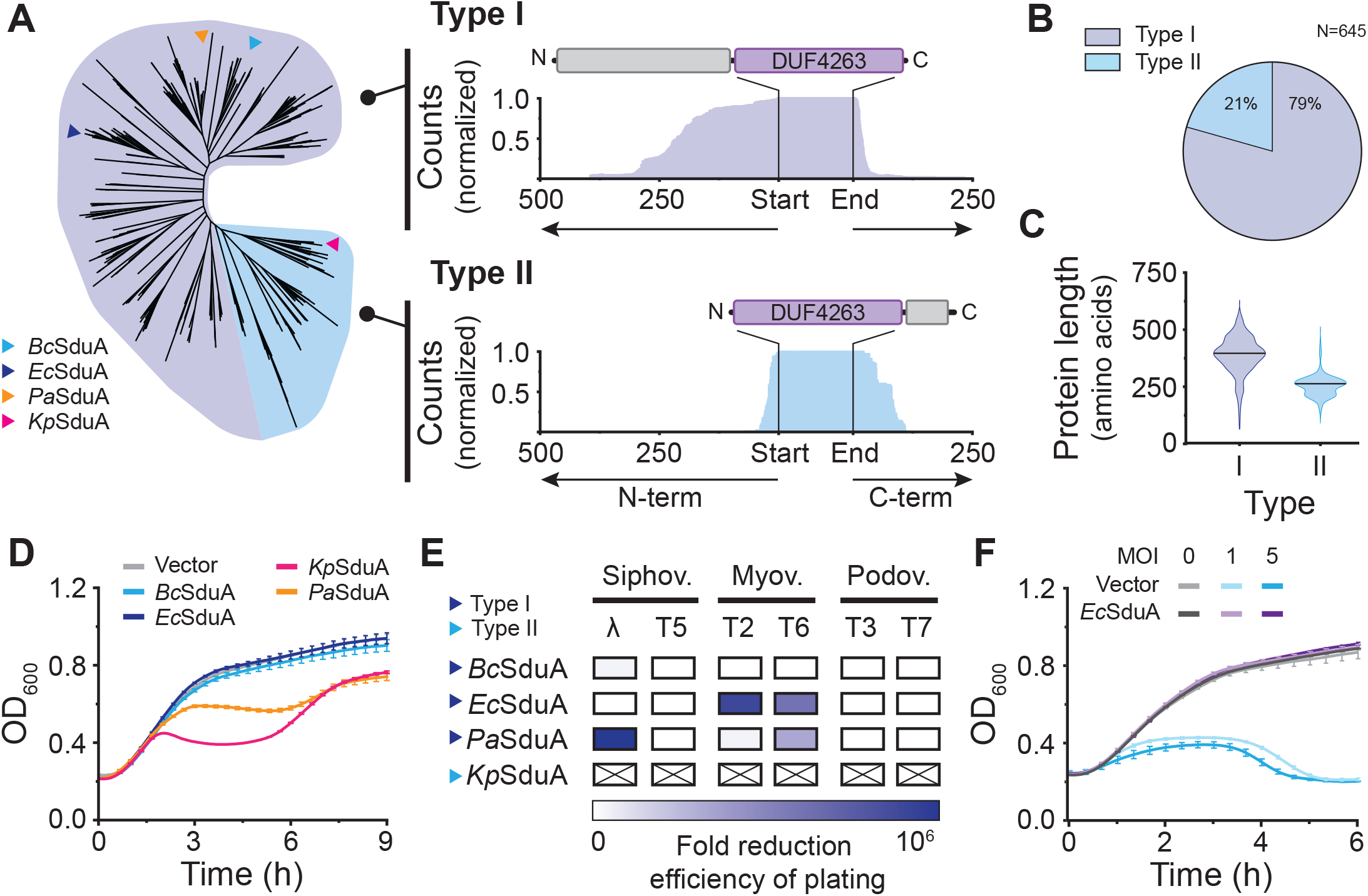
Architecture of Shedu anti-phage defense systems. **(A)** Phylogenic classification of 645 Shedu systems into two distinct subtypes, based on the domain architecture of SduA. In type I systems (purple) the hallmark domain DUF4262 is positioned at the C-terminus of the protein with a large N-terminal region. In type II systems (light blue) DUF4262 is flanked by short N- or C-terminal regions. **(B)** Prevalence of type I (purple) and type II (light blue) systems in percent. **(C)** Violin plots showing the polypeptide length distribution of type I (purple) and type II (light blue) Shedu systems. Solid black line indicates the median length. **(D)** Growth curves of *E. coli* cells expressing Shedu orthologs. Negative control strain harbors empty expression vector. Data points represent mean ± SEM of three biological replicates (n=3). **(E)** Anti-phage defense phenotype of *E. coli* cells expressing Shedu orthologs against six *E. coli* phages. Color intensity represents fold reduction in the efficiency of plating (EOP), also see Figure S1D. KpSduA displayed intrinsic cellular toxicity. **(F)** Growth curves for *E. coli* cells harboring the EcSduA system or an empty vector, infected with the T6 phage at a multiplicity of infection (MOI) of 0, 1 or 5. Data points represent mean ± SEM of three biological replicates (n=3).

### Shedu provides anti-phage immunity

To understand the function of Shedu *in vivo*, we expressed distinct orthologs of Shedu in *E. coli* cells using an arabinose inducible promoter and tested their effect on cell growth. Growth curves showed that cells expressing Shedu orthologs from *E. coli* KTE10 (EcSduA, type I) or *Bacillus cereus* B4262 (BcSduA, type I) grew at similar rates compared to cells harboring an empty expression vector (**Figure 1D**). By contrast, cells expressing SduA orthologs from *Pseudomonas aeruginosa* O12 (PaSduA, type I) and *Klebsiella pneumoniae* BIDMC47 (KpSduA, type II) grew at slower rates compared to the empty-vector control, suggesting that these orthologs cause cellular toxicity when expressed in a non-native host (**Figure 1D**). This observation suggests that the activities of Shedu systems have been adapted to their hosts and might provide phage defense in a host-specific manner.

Next, we challenged *E. coli* strains harboring each Shedu system with a set of six of *E. coli* phages that belong to the three major families of tailed dsDNA phages (*Sipho-, Myo-*, and *Podoviridae*). The EcSduA ortholog provided a >30,000-fold reduction in the efficiency of plaquing (EOP) against *Myoviridae* bacteriophages T2 and T6, as compared to the vector-only control, indicative of a strong defense phenoype (**Figure 1E** and **S1D**). PaSduA showed a strong defense phenotype against lambda phage, >1,000,000-fold EOP reduction, and >80-fold EOP reduction when challenged with T6, whereas only a mild defense phenotype was observed against T2 with a 2.9-fold EOP reduction. When we tested the BcSduA ortholog for phage defense in *E. coli*, which was previously shown to provide phage defense in *Bacillus subtilis*^2^, we observed a mild (<7.5-fold) reduction in the EOP against bacteriophage lambda. These observations highlight that Shedu systems provide phage-specific host defense and have evolved to target specific phages that infect their hosts.

Given that EcSduA exhibited a strong defense phenotype against a single family of phages, we decided to use EcSduA as a model system to elucidate the molecular mechanisms that underpin the immune function of Shedu systems. As over 70% of prokaryotic gene defense systems characterized to date mediate immunity by causing programmed cell death to abort phage infection^27,28^, we assessed whether EcSduA provides phage defense through an abortive infection mechanism. When cells lacking EcSduA were challenged with T6 phage at a low and high MOI in liquid culture, cell cultures collapsed due to the T6 lytic replication cycle. In contrast, cells harboring EcSduA resisted infection even at a high MOI and grew at the same rate as the uninfected control (**Figure 1F**), indicating that Shedu does not elicit programmed cell death upon phage infection. This suggests that Shedu instead provides immunity through a mechanism that inhibits phage replication, and implies that Shedu systems are capable of self versus non-self discrimination.

### Molecular architecture of Shedu

To understand the defense mechanism of Shedu, we sought to obtain structural insights into the EcSduA protein. Recombinant expression and purification of EcSduA indicated that the protein formed oligomers as indicated by size exclusion chromatography (**Figure S2A**). Subsequent analysis by size exclusion chromatography coupled with multiple angle light scattering (SEC-MALS) revealed a mono-disperse peak with a total mass of 194.8 ± 1.8 kDa, corresponding to a tetrameric EcSduA complex (**Figure S2B**). We then subjected EcSduA to structural analysis by single-particle cryo-EM, obtaining a reconstruction with a nominal resolution of 3.0 Å (**Figure 2A** and **S3A-C**). To facilitate building of an atomic model, we additionally determined a crystal structure of the EcSduA N-terminal domain at a resolution of 2.5 Å (**Figure S2D**). The cryo-EM structure of EcSduA reveals a C2-symmetric, dimer-of-dimers assembly, in which individual protomers (SduA.1/SduA.3 and SduA.2/SduA.4) dimerize through an extensive hydrophobic interface that spans a surface area of ∼4100 Å^2^ (**Figure 2B** and **S4A-C**). The two SduA dimers assemble into the tetramer via the DUF4263 domains of protomers SduA.2 and SduA.4, forming an interface with a surface area of ∼1450 Å^2^ (**Figure 2B** and **S4A-C**). As a result, the central body of the tetrameric complex is formed by the DUF4263 domains, while the N-terminal domains project on opposite sides of the central body, forming two clamp-like structures (**Figure 2B**).

**Figure 2:**
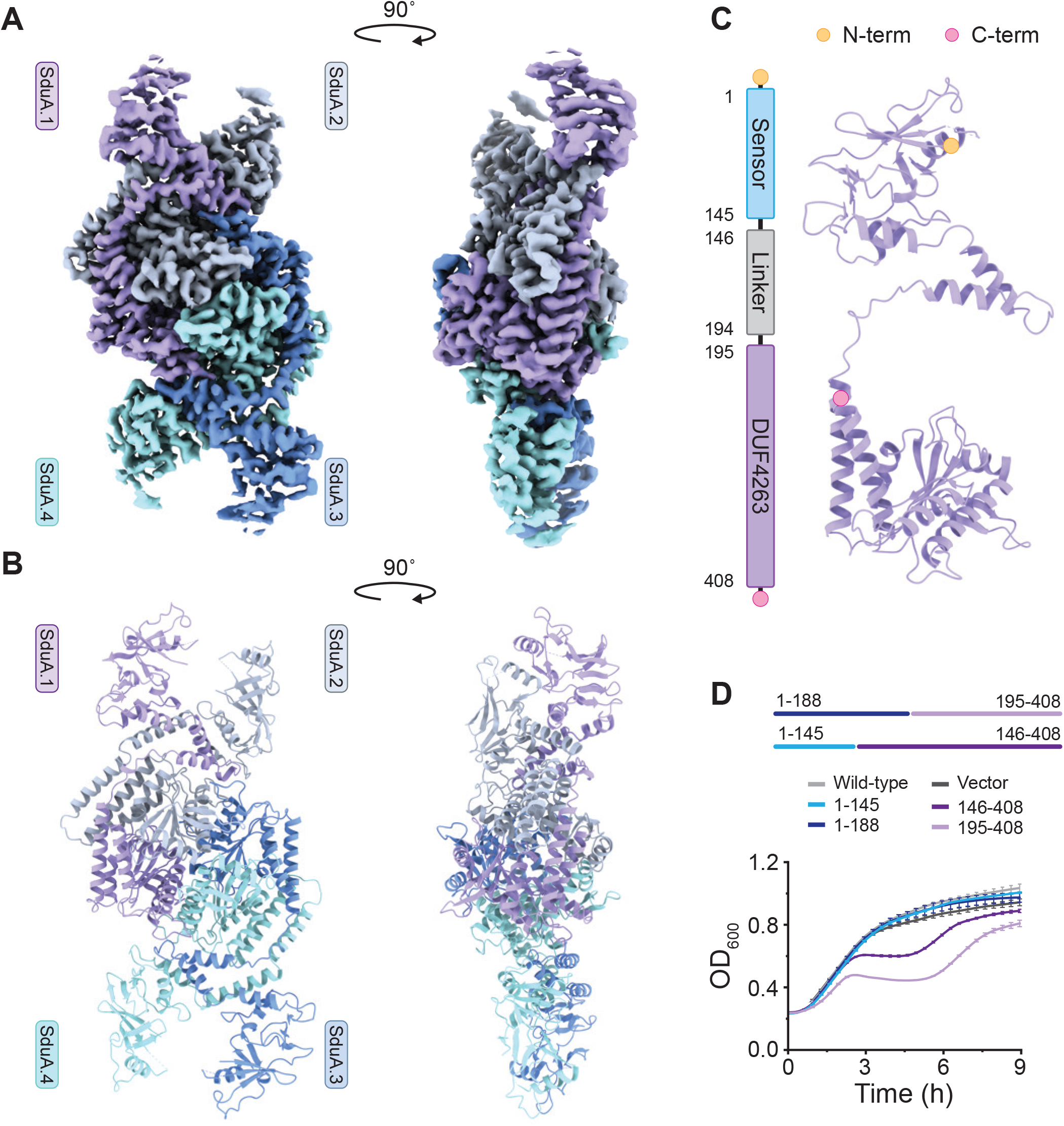
Structure of the Shedu anti-phage defense complex. **(A)** Cryo-EM density map of the tetrameric EcSduA complex. **(B)** Structural model of tetrameric EcSduA, shown in cartoon representation. **(C)** Structural model of the SduA.1 monomer (right) shown alongside domain organisation (right). N- and C-termini are indicated by orange and pink circles, respectively. **(D)** Growth curves for *E. coli* cells expressing various truncations of EcSduA. Curves represent mean ± SEM of three biological replicates (n=3).

Each SduA protomer is composed of two domains connected by an alpha helical linker: an N-terminal domain that lacks structural homology to other proteins and a C-terminal DUF4263 nuclease domain (**Figure 2C** and **S4D**). To understand the functional roles of these domains, we expressed truncated forms of EcSduA in *E. coli* and monitored cell growth. When the N-terminal domain (residues 1-145) was expressed alone or fused with the alpha-helical linker (residues 1-188), cells grew at similar rates as those expressing full-length EcSduA or vector-only control cells (**Figure 2D**). In contrast, expression of the signature DUF4263 domain (residues 195-408) alone or as a fusion with the alpha-helical linker (residues 146-408), resulted in cell toxicity and growth inhibition (**Figure 2D**). These data indicate that the N-terminal domain regulates the activity of the DUF4263 domain, and suggest that the N-terminal domain may acts as a sensor that provides target specificity to the Shedu system.

### Shedu is a dsDNA-targeting nuclease

Given the requirement of the DUF4263 domain for the interference function of type I Shedu immune systems, we analyzed the nuclease activity of EcSduA *in vitro* using 5’-fluorophore-labeled synthetic oligonucleotide substrates. Upon incubating EcSduA with a ssDNA or single-stranded RNA (ssRNA) substrates, no cleavage was observed (**Figure 3A**). By contrast, incubation of dsDNA in the presence of EcSduA and Mg^2+^ resulted in the accumulation of a degradation product of approximately 10-15 nucleotides in length, suggesting that EcSduA is a dsDNA targeting system (**Figure 3A**). In agreement with this observation, a fluorescence polarization binding assay carried out in the absence of divalent ions showed that EcSduA has high affinity for dsDNA, displaying an apparent dissociation constant (*K*_D_) of 11.9 ± 0.4 nM (**Figure 3B**). Taken together these data suggest that Shedu provides immunity by directly targeting dsDNA of invading phages.

**Figure 3:**
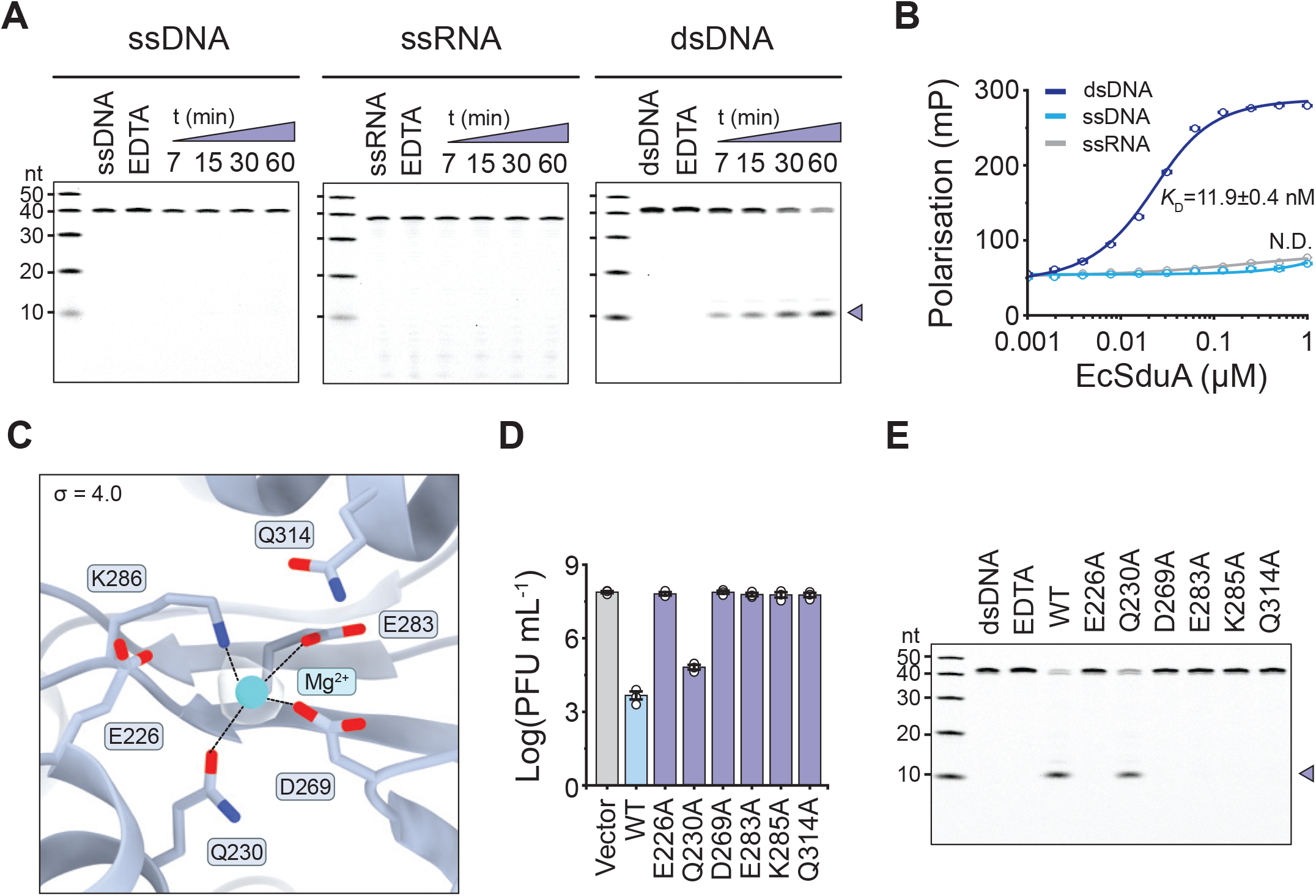
Shedu cleaves dsDNA substrates using a PD/ExK nuclease domain. **(A)** *In vitro* nuclease activity assays using fluorescently labelled oligonucleotides. Cleavage product (indicated with purple triangle) was resolved by denaturing polyacrylamide gel electrophoresis (PAGE). **(B)** Binding curves of EcSduA to fluorescently labelled oligo nucleotides measured by fluorescence polarization. Data represent the mean ± SEM of three independent replicates (n=3). Solid line represents a nonlinear regression fit. **(C)** Close-up view of the active site of EcSduA showing catalytic residues that coordinate a magnesium ion (depicted as cyan sphere). Map (light grey) contoured at σ=4. **(D)** Plaque assays of T6 phage infecting *E. coli* strains expressing wild-type (WT) EcSduA, EcSduA active site mutants or a vector-only control. Data represents mean PFU mL^-1^ ± SEM of three independent replicates (n=3). **(E)** Nuclease activity on a fluorescently labelled dsDNA oligonucleotide by WT EcSduA and active site mutants. Cleavage product (indicated with purple triangle) was resolved by denaturing PAGE.

The EcSduA structure reveals that the DUF4262 domain comprises a central five-stranded beta sheet decorated by alpha helices (**Figure 2C** and **S4D**), which resembles the signature core of proteins belonging to the PD-(D/E)xK phosphodiesterase superfamily_^29^_ (**Figure S4E**). A conserved active site pocket lined with residues Glu226, Gln230, Asp269, Glu283, Lys285 an Gln314 features density corresponding to a coordinated magnesium ion in the cryo-EM reconstruction (**Figure 3C** and **S4F-G**). In agreement with these observations, alanine substitutions of Glu226, Asp269, Glu283, Lys285 and Glu314 abolished SduA-mediated anti-phage immunity in *E. coli* and inhibited dsDNA cleavage *in vitro*, while mutation of Gln230 resulted in a reduced immunity *in vivo* (**Figure 3D-E** and **S4H**). Taken together these results confirm that SduA is a dsDNA-selective PD-(D/E)xK-like nuclease whose catalytic activity is essential for phage defense by the Shedu immune system.

### Structural basis for dsDNA sensing by Shedu

We next focused on the mechanism of phage sensing and self-versus-non-self discrimination. The observation that the N-terminal domain of EcSduA is necessary to limit the toxicity of the DUF4263 nuclease domain *in vivo* suggested that the N-terminal domain may act as a sensor responsible for target recognition in Shedu systems. Given the strong phage defense phenotype of EcSduA against T-even phages of the *Myoviridae* family (**Figure 1E**), we initially hypothesized that EcSduA may recognize the glucosyl modification of 5-hydroxymethylated cytosines found in the DNA of T-even phages^30,31^. However, neither the glucosylated genomic DNA from T2 and T6 phages, nor unmodified genomic phage DNA from T7 showed substantial degradation *in vitro*, indicating EcSduA does not specifically recognize glycosylated dsDNA substrates (**Figure S5A**). This observation contrasts with robust cleavage of short dsDNA oligonucleotides *in vitro* (**Figure 3A**), suggesting that EcSduA may instead target dsDNA intermediates that occur during phage replication.

Based on the strong affinity of EcSduA for short dsDNA oligonucleotides, we reconstituted a complex of EcSduA with a 40-base pair (bp) palindromic dsDNA oligonucleotide in the absence of divalent metal ions, and determined its structure by single-particle cryo-EM at a nominal resolution of 2.8 Å (**Figure 4A** and **S6A-C**). The dsDNA is bound along one face of the EcSduA tetramer, spanning its entire length. Even though no symmetry was imposed during cryo-EM data processing, the resulting reconstruction exhibits pseudo-C2 symmetry and the bound dsDNA in the final reconstruction appears to be longer than 40-bp, likely as a result of averaging during particle alignment. Strikingly, the EcSduA-dsDNA complex structure reveals that the end of the dsDNA substrate is inserted into the clamp-like structure formed by the N-terminal domains of protomers SduA.1 and SduA.2 (**Figure 4B** and **4C**). To accommodate the ∼24 Å wide B-form DNA in the 17 Å wide N-terminal domain clamp, the bound dsDNA duplex is unwound at the end while preserving its base-pairing (**Figure 4D, S5B** and **S5D**). Consistent with this, modeling of an extended B-form DNA duplex in the N-terminal domain clamp of the EcSduA results in steric clashes with one of the protomers (SduA.1), suggesting that the N-terminal domain clamp is only able to bind to ends of linear dsDNA molecules (**Figure S5C**). The bound DNA end is stabilized by a highly conserved Trp25 residue protruding from the N-terminal domain of protomer SduA.2; in accordance, alanine substitution of this residue abolished phage resistance *in vivo* (**Figure 4E-F)**.

**Figure 4:**
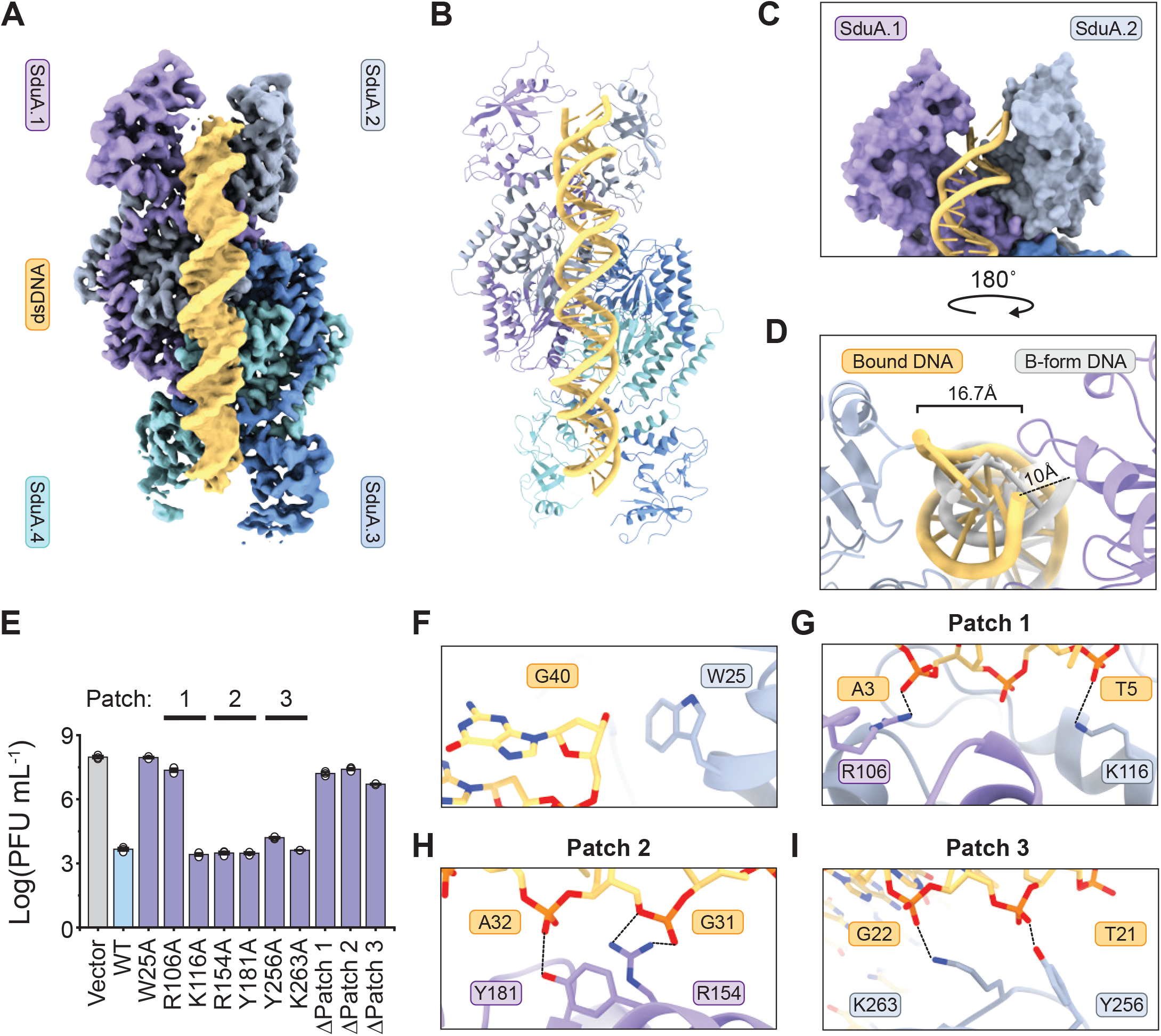
Shedu senses dsDNA termini. **(A)** Cryo-EM density map of tetrameric EcSduA in complex with a dsDNA substrate. **(B)** Atomic model of the EcSduA-dsDNA complex. **(C)** Close-up view of the N-terminal domains of EcSduA bound to the end of the dsDNA substrate. **(D)** Structural superposition of the dsDNA substrate bound by the EcSduA complex (light orange) and an ideal B-form dsDNA (light gray). **(E)** Plaque assay of T6 phage infecting *E. coli* strains expressing WT EcSduA, EcSduA substrate binding mutants or a vector-only control. Data represents the mean PFU mL^-1^ ± SEM of three independent replicates (n=3). **(F)** Detailed view of the interactions at the end of the dsDNA substrate. **(G)** Detailed view of the interactions with the dsDNA backbone within Patch 1. **(H)** Detailed view of the interactions with the dsDNA backbone within Patch 2. **(I)** Detailed view of the interactions with the dsDNA backbone within Patch 3.

From the N-terminal clamp, the dsDNA extends over the center of the EcSduA tetramer, positioned by a positively charged binding channel (**Figure S5E**), which introduces a ∼24° convex bend in the dsDNA (**Figure S5F**). The negatively charged dsDNA backbone is engaged in extensive electrostatic and hydrogen bonding contacts with three distinct interaction interfaces (referred to as patch 1 to 3) in EcSduA (**Figure 4G-I** and **S5E**). Apart from these interactions with the DNA backbone, no base-specific interactions were observed, indicating that EcSduA does not bind dsDNA in a sequence-specific manner. In agreement with the observed mode of dsDNA binding, alanine substitution of Arg106 within patch 1 was sufficient to abolish immunity *in vivo*. Although other individual mutations were not sufficient to perturb activity, combined alanine substitutions of residues within patch 1 (Arg106 and Lys116), patch 2 (Arg154 and Tyr181), or patch 3 (Tyr256 and Lys263) resulted in a complete loss of phage resistance (**Figure 4E)**. Together, these results validate the observed mode of dsDNA binding by EcSduA and suggest that the Shedu system recognizes ends of linear dsDNA targets.

### Shedu nicks linear dsDNA at fixed distance from 5’ ends

The terameric EcSduA assembly features four DUF4263 nuclease domain active sites, two on each face of the tetramer. The structure of the EcSduA-dsDNA complex reveals that upon end binding by the N-terminal domain clamp, the dsDNA substrate projects over the nuclease active site of the DU4263 domain protomer SduA.2 and extends towards the active site in symmetry-related protomer SduA.4. The backbone of the 5’-strand is positioned such that the phosphate of nucleotide 14 points towards the SduA.2 catalytic center. These findings suggest that EcSduA nicks dsDNA substrates on the 5’ strand, at a fixed distance downstream from the 5’ end (**Figure 5A**), in agreement with the observation of a ∼13-nt cleavage product in nuclease activity assays (**Figure 3A** and **S5G**). Notably, the backbone of the substrate DNA strand is positioned ∼12 Å away from the catalytic center, too far away for catalysis even when accounting for the absence of divalent metal ions, suggesting that the DNA substrate and/ or EcSduA undergo conformational rearrangements upon activation by divalent metal ions.

**Figure 5:**
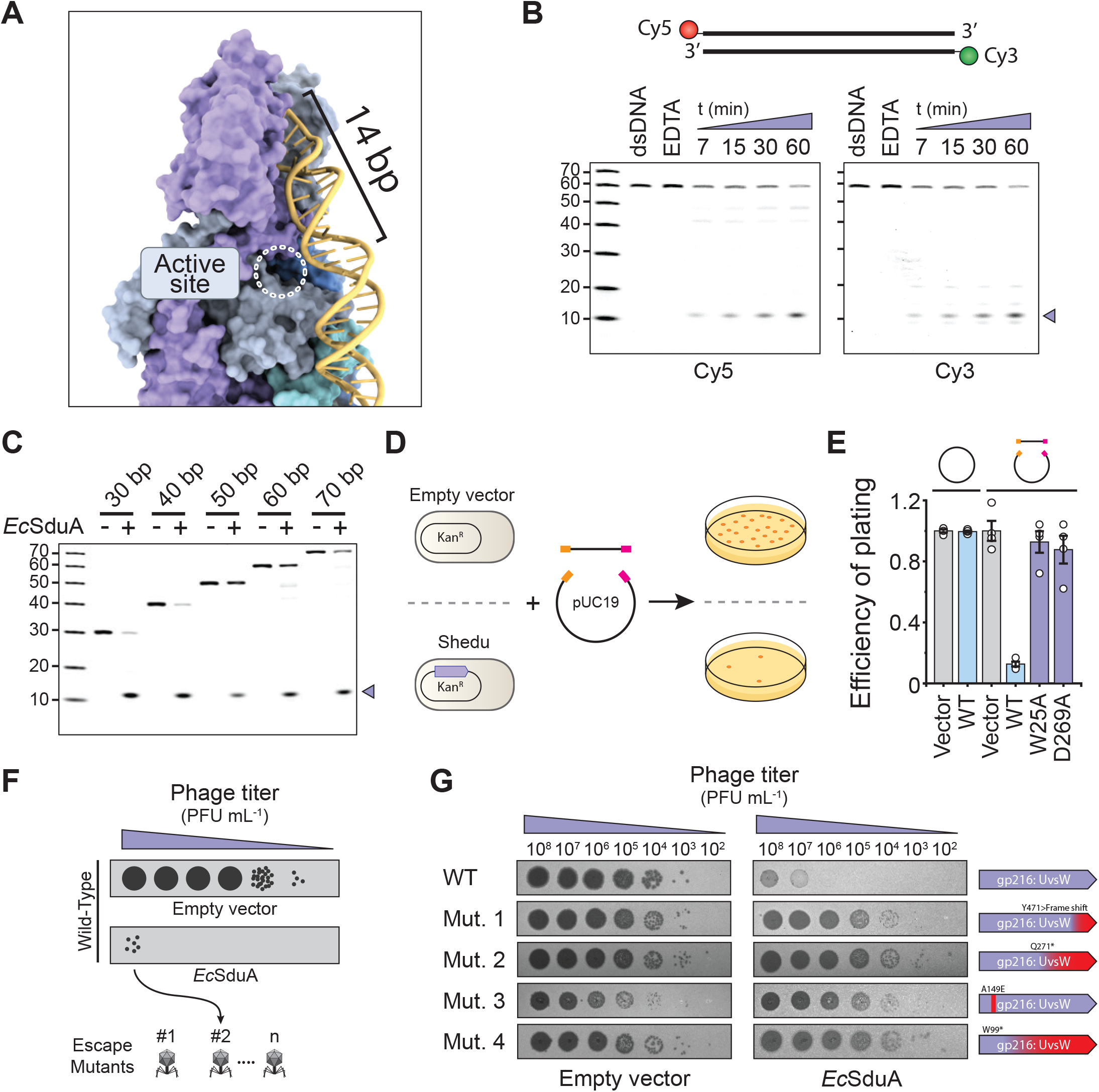
Mechanism of dsDNA cleavage and phage immune evasion. **(A)** Detailed view of the path of the dsDNA substrate from the end-binding site over the active site. **(B)** Nickase activity of WT EcSduA on a doubly-labelled dsDNA oligonucleotide. Nicking products (indicated with purple triangle) were resolved by denaturing PAGE. **(C)** Analysis of nicking activity of WT EcSduA on fluorescently labelled dsDNA oligonucleotides of increasing lengths. Nicking products (indicated with purple triangle) were resolved by denaturing PAGE. **(D)** Schematic showing an *in vivo* DNA transformation assay to probe end recognition by the Shedu immune system. *E. coli* strains expressing WT or mutant EcSduA proteins (or vector-only control strain) were transformed with circular plasmid or linear dsDNA fragments, after which transformants were counted. **(E)** Normalized transformation efficiency of circular plasmid or linear dsDNA fragments in presence and absence of WT or mutant EcSduA. Data represents mean transformation efficiency of three independent replicates ± SEM (n=3). Data was normalized against the vector-only control for the respective substrate. **(F)** Schematic depiction of T6 escaper phage isolation procedure. **(G)** Left: Analysis of the escape phenotype of the isolated phages by serial-dilution plaque assay (left). Right: Depiction of mutations in the *uvsW* gene in isolated T6 escaper phages, including frame shifts and pre-mature stop codons, as determined by whole genome sequencing.

If Shedu specifically recognizes dsDNA ends and nicks at a fixed distance downstream from the 5’ end, we reasoned that both ends of a linear dsDNA oligonucleotide would exhibit the same cleavage pattern *in vitro*. In agreement with this hypothesis, nuclease activity assays with a doubly labelled DNA substrate revealed that both ends were nicked at similar positions, based on the apparent sizes of the cleavage products (**Figure 5B**). Likewise, when we tested a set of dsDNA substrates that varied in length, we observed the same main ∼13 nt cleavage product for all substrates (**Figure 5C** and **S5G**). For substrates that were longer than 50 bp, we observed additional minor cleavage products, possibly arising from cleavage by the active site of protomer SduA.4 in the distal half of the tetramer (**Figure 5C**).

To validate dsDNA end recognition and cleavage mechanisms of EcSduA *in vivo*, we designed a plasmid-based transformation assay that probes the activity of Shedu independent of phage infection (**Figure 5D**). To this end, we generated chemically competent *E. coli* strains carrying an expression plasmid for wild-type (wt) EcSduA or an empty control plasmid. Next, these strains were transformed with either an intact circular plasmid carrying an ampicillin resistance marker or two linear plasmid-derived fragments carrying 15-bp homology arms (**Figure 5D**). When cells were transformed with the intact plasmid, both the strain expressing wt EcSduA or the control strain yielded comparable numbers of transformants, indicating that Shedu does not target plasmids. By contrast, when the assay was repeated with DNA fragments, the strain expressing wt EcSduA yielded ∼10-fold fewer transformants as compared to the empty-vector control strain (**Figure 5D**). Strains expressing EcSduA variants with mutations in the sensor or nuclease domains that abolish phage defense *in vivo* (W25A or D269A) yielded comparable numbers of transformants as the vector-only control strain (**Figure 5D**). Taken together, these results confirm that Shedu specifically recognizes and nicks linear dsDNA substrates at their termini, both *in vitro* and *in vivo*.

### Phages evade Shedu by suppressing linear DNA replication intermediates

To provide further evidence for the mechanism of the Shedu immune system, we sought to isolate Shedu-resistant phages and identify the genetic mutations underpinning their immune evasion. To this end, a culture of *E. coli* expressing WT EcSduA was exposed to T6 phage, after which surviving plaques were isolated and amplified. This procedure yielded seven T6 mutant escaper isolates that resisted host immunity (**Figure 5F**). Whole-genome sequencing revealed that the isolated phages contained multiple mutations, including point mutations as well as large deletions up to several kilobases (**Figure S7A**). Close inspection of the mutations across the escaper strains revealed that most mutations were not conserved except for missense and nonsense mutations in gene product 216, which encodes the ATP-dependent DNA helicase UvsW (**Figure 5F** and **Figure S7B**). UvsW functions together with recombination mediator proteins UvsX and UvsY and drives Holliday junction branch migration in both homology directed DNA repair and recombination-dependent replication pathway in T-even phages^32–34^. The loss of UvsW thus might explain the accumulation of spurious mutations and large genomic deletions in escaper phages (**Figure S7A**). As the recombination-dependent replication pathway of T-even phages relies on the generation of linear dsDNA intermediates that serve as templates for homologous recombination, the loss of UvsW in escaper phages is consistent with a functional mechanism of Shedu based on recognition and nicking of linear dsDNA molecules.

## Discussion

Innate immune systems detect pathogen-associated molecular patterns (PAMPs) of invaders to rapidly counter the incoming infection^35–38^. To make effective targets for innate immune systems, PAMPs must share three common features^39^. First, to avoid auto immunity by the innate immune system, the PAMP should only be present in the pathogen and not in the host. Second, to provide protection against a broad spectrum of invaders, the recognized PAMP should have an invariant feature that is conserved among a class of pathogens. Third, the PAMPs should be essential for the survival of the pathogen, preventing the emergence of mutants that escape the innate immune system.

The constant evolutionary arms race between phages and their prokaryotic hosts has resulted in a myriad of innate defense mechanisms that provide robust immunity through distinct enzymatic activities. In this study, we employed an integrative experimental approach to reveal the molecular mechanism of the Shedu anti-phage defense system. Our findings suggest that a subset of Shedu systems use the N-terminal domain of SduA to recognize free ends of linear DNA molecules, thus providing a mechanism for phage sensing. Subsequently, SduA nicks the DNA using its hallmark DUF4264 PD/ ExK-like nuclease domain, thereby impairing phage proliferation (**Figure 6**). The model is consistent with the observations that Shedu does not target circular plasmid molecules *in vivo*; in turn, the genomes of the T-even phages targeted by the EcSduA exist in the linear form upon host infection and undergo replication by a recombination-based mechanism that involves linear DNA segments^40^. Together, these results underscore that the Shedu innate immune system functions through molecular pattern recognition by sensing a phage-specific PAMP that is absent from the host.

**Figure 6:**
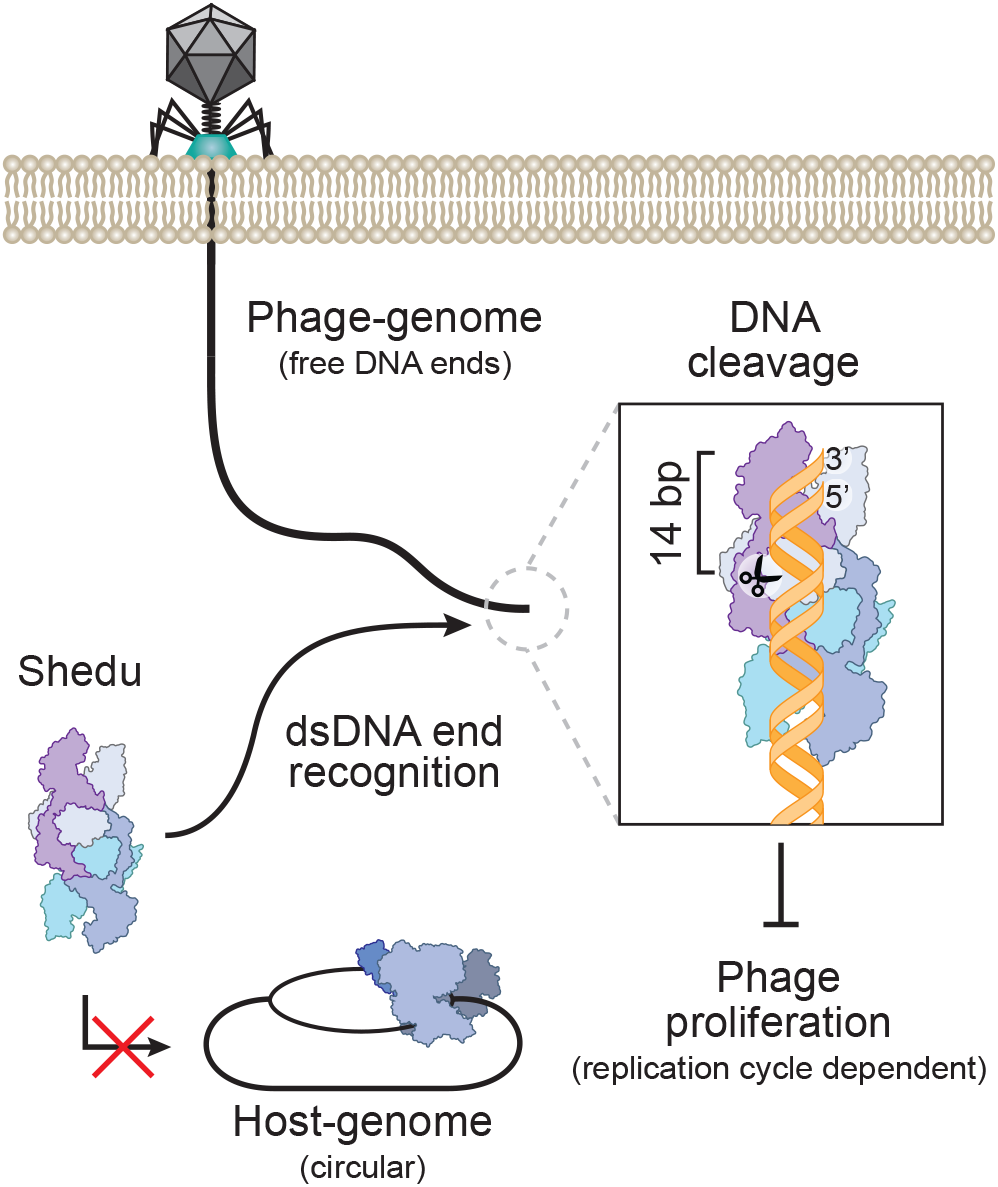
Pattern recognition by the Shedu immune system. Model for the anti-phage activity of the Shedu immune system. Shedu recognizes free ends of phage genomic DNA upon infection and/or recombination-dependent replication. Recognition of the dsDNA termini results in nicking 14-bp away from the 5’ end. Depending on the replicative cycle of the invading phage, the end cleavage impairs phage proliferation. The host genome remains circular throughout its replicative cycle and is thereby unaffected by the Shedu immune system.

Consistent with the initial study of Shedu^2^, our functional analysis of several Shedu orthologs shows that the systems provide bacteriophage-specific immunity (**Figure 1E**), suggesting that Shedu systems are tailored to the phages that infect their hosts. Our bioinformatic analysis shows that the N-terminal domains of type I SduA proteins are highly variable, suggesting that the N-terminal sensor domains are evolutionarily tailored to recognize specific phage families and implying that some Shedu systems might recognize phage-specific molecular signatures other than free DNA ends. The molecular mechanism of type II Shedu systems, in which the SduA protein lacks any discernible domains beyond the DU4263 nuclease domain, is presently unclear. It is likely that these systems employ additional, possibly host-encoded, co-factors for target recognition. The presence of Shedu systems in Z/fun locus, a hotspot for anti-phage defense systems^41^ in P2 prophages (**Figure S1E**), further highlights that Shedu provides immunity to specific phage families and suggests that Shedu systems might additionally participate in the competition of mobile genetic elements for their host.

Whole genome sequencing of mutant phages that evade EcShedu indicates that they escape through a mutation in the *uvsW* gene. UvsW is a branch-migrating dsDNA helicase that plays a critical role in the recombination-dependent replication (RDR) cycle of T-even phages^32–34^. Upon host infection, the initial stage of the replication the genome of T-even phages relies almost completely on the RDR pathway, in which proteins UvsY and UvsX catalyze recombination-mediated strand invasion of the ends of linear DNA intermediates^40,42^. The resulting Holliday junction intermediates are subsequently resolved by UvsW through branch-migration, which is followed by synthesis of new strands by DNA polymerase^40^. As DNA ends serve as substrates for SduA (**Figure 6**), this implies that Shedu targets DNA replication intermediates, thereby impairing phage proliferation upon infection. Loss of UvsW in T-even phages has been shown to facilitate a switch to an alternative mode of replication^32^, thus suggesting an explanation for the evasion of Shedu immunity by *uvsW* phage mutants (**Figure 6**).

Many bacteria, including *E. coli*, encode the RecBCD machinery that plays a dual role in dsDNA break repair and anti-viral immunity^15,43,44^. RecBCD is a potent dsDNA-specific helicase-exonuclease that targets linear products generated by restriction of foreign DNA^43,45^. To counteract the function of RecBCD, phages encode potent inhibitors that prevent dsDNA targeting by the RecBCD machinery through distinctive strategies^15,46,47^. The redundancy in targeting DNA ends by RecBCD and Shedu systems is consistent with the pan-immune system theory^7^, in which phage defense systems cooperate as a bet-hedging strategy to ensure host survival. In analogy with anti-RecBCD proteins that mimic DNA substrates^46,47^, we anticipate that phages encode anti-Shedu proteins to evade Shedu immunity.

The dsDNA end-sensing mechanism of Shedu raises parallels with eukaryotic innate immune pathways that sense nucleic acid duplex ends as PAMPs. The vertebrate cytosolic pattern recognition receptor RIG-I specifically recognizes the termini of short 5’-triphosphorylated dsRNA substrates to trigger the downstream type-1 interferon innate immune response^48–50^. The *Drosophila melanogaster* Dicer-2 RNase III enzyme recognizes the termini of dsRNA substrates to generate siRNA guides for Argonaute-mediated silencing of invading viruses^51–54^. The discovery of a prokaryotic antiviral system that recognizes and targets dsDNA ends thus underscores the paradigm that recognition of pathogen-specific nucleic acid structures is a conserved feature of innate immunity across the domains of life.

### Limitations of the study

Our findings point towards a mechanism in which the N-terminal domains of SduA sense free ends of dsDNA. Yet, the N-terminal domains SduA proteins are highly diverse across Shedu systems, suggesting that they may provide other, distinct mechanisms of phage detection. A major focus of future experiments will be to understand how these distinct protein architectures provide phage-specific targeting.

Our structural analysis of the EcSduA complex bound to dsDNA shows that the substrate runs over an active site. Yet, the distance between the DNA backbone and the active site is too far for catalysis, implying that a conformational rearrangement in the SduA protein or the bound DNA is necessary to enable DNA nicking. The underlying mechanistic details of catalytic activation remain to be determined. Notably, the density corresponding to the bound duplex DNA in the cryo-EM structure of the EcSduA-dsDNA complex is larger than expected; furthermore, the DNA extends from the N-terminal domain clamp at one end of the tetrameric complex towards the N-terminal clamp on the opposite side. The artifactual length of the bound DNA and the pseudosymmetry of the complex is possibly due to particle misalignment and averaging during cryo-EM data processing.

## Acknowledgements

We are grateful to Marta Sawicka, Simona Sorrentino and Piotr Szwedziak (University of Zurich Center for Microscopy and Image Analysis) for technical support with cryo-EM data acquisition. We thank Weihong Qi and Maria Domenica Moccia from the Functional Genomics Center Zurich for assistance with deep sequencing. We thank Meitiwan Wang, Takashi Tomizaki and Vincent Olieric (Paul Scherrer Institute) for assistance with X-ray diffraction data collection. We thank Kevin Corbett and Yajie Gu for fruitful discussions and sharing unpublished data. We thank all the members of the Jinek lab for constructive feedback throughout the project. The work was funded by a European Research Council Consolidator Grant (project no. 820152, CRISPR2.0) to MJ. LL was funded by a Horizon 2020 Marie Skłodowska-Curie Individual Fellowship (project no. 845268, MSOPGDM). MJ is International Research Scholar of the Howard Hughes Medical Institute, Vallee Scholar of the Vallee Foundation and member of the Swiss National Competence Center for Research “RNA & Disease”.

## Author Contributions

LL and MJ conceived study. LL performed bioinformatic analysis, biochemical experiments, phage infection experiments and *in vivo* assays, and analyzed deep-sequencing data. LL, AW and GTR purified proteins. LL prepared cryo-EM samples, collected and analyzed cryo-EM data. LL and AW crystallized proteins and collected X-ray diffraction data. LL and MJ wrote manuscript with input from all authors.

## Resource availability

### Lead contact

Further information and requests for resources and reagents should be directed to and will be fulfilled by the Lead Contact, Martin Jinek.

### Materials availability

Proprietary material is available upon request from the authors.

### Data and code availability

Atomic coordinates and cryo-EM maps for the apo EcSduA complex (PDB: 8PS4), EMDB: EMD-17844) and the DNA-bound EcSduA complex (PDB: 8PS5), EMDB: EMD-17845) have been deposited in the PDB and EMDB databases. Structure factors and atomic coordinates of the reported X-ray crystallographic structure of the N-terminal fragment of EcSduA has been deposited in the Protein Data Bank with accession code 8PS6 Accession codes are listed in the key resources table.

## Methods

### Bioinformatic analysis of Shedu orthologs

Bioinformatic analysis was performed on the list of SduA orthologs that was published by Doron et al.^2^. Redundant protein coding sequences were removed from the list using a custom written MATLAB script (Mathworks), resulting in a list of 645 unique SduA orthologs. The sequences of these orthologs were retrieved from NCBI and aligned using MAFFT (v7.490)^57^, after which a phylogenetic tree was generated using IQ-tree (v2.1.3)^58^ with automated model selection^59^ and visualized using FigTree (version 1.4.4). For the classification of the two distinct subtypes, the domain boundaries of DUF4263 were determined with HHMER (v3.3.2)^60^, after which the number of amino acids preceding and succeeding the DUF4263 was analyzed determined for each ortholog using a custom written MATLAB script (Mathworks).

### Plasmid DNA constructs and site-specific mutants

The DNA sequences of *Bacillus cereus* SduA (ACK61957.1), *Escherichia coli* SduA (ELC07825.1), *Klebsiella pneumoniae* SduA (EWE02243.1) and *Pseudomonas aeruginosa* SduA (KQJ67128.1) were codon optimized for heterologous expression in *E. coli* and synthesized by GeneArt (Thermo Fisher Scientific). The EcSduA gene was inserted into the 1B (Addgene: 29653) plasmid using ligation-independent cloning (LIC), resulting in a construct that carries an N-terminal hexahistidine tag followed by a tobacco etch virus (TEV) protease cleavage site. The genes of BcSduA, EcSduA and truncated derivatives and mutants, KpSduA and PaSduA were inserted into the 8A (Addgene: 37501) plasmid using ligation-independent cloning (LIC), resulting in a tag free construct that is under control of an arabinose inducible promotor. Site specific mutations were introduced by QuickChange mutagenesis or by inverse PCR. Plasmids were purified using the GeneJET plasmid miniprep kit (Thermo Fisher Scientific) and insertion and mutagenesis were verified by Sanger sequencing.

### Phage cultivation and plaque assays

Phage experiments with phages T2 (DSM16352), T3 (DSM4621), T5 (DSM16353), T6 (DSM4622) and T7 (DSM4623) were performed in *E. coli* strain DSM613, whereas experiments with Lambda phage (DSM4499) were performed in *E. coli* strain K12 (DSM4230). To generate phage stocks, *E. coli* cells were grown in lysogeny broth (LB) supplemented with 1 mM MgCl_2_, at 37 °C, 220 rpm until reaching an optical density at 600 nm (OD_600_) of 0.3. Cultures were subsequently infected with phage and incubation was continued until collapse of the culture. To clean up the lysates, 1% cholorophorm was added together with 10 μg mL^-1^ DNase 1 (NEB) and 1 μg mL^-1^ RNase A, followed by incubation for 1 hour at 37°C. After incubation, samples were cleared by centrifugation at 15,000g for 10 minutes, supplemented with NaCl to 0.5M and stored at 4°C until the titer was determined with a plaque assay.

For plaque assays the respective *E. coli* strain was grown in LB until reaching an OD_600_ of 1.0. Next, 400 cells μL were mixed with 50 μL diluted phage stock and 4 mL lukewarm top agar (LB supplemented with 7 g L^-1^ agar and 0.1% L-arabinose) and poured onto 9 cm LB agar plate with 0.1% L-arabinose at room temperature. Plates were incubated overnight at 37°C and the next day plaque forming units (PFU) were counted and used to calculate the viral titer. Stocks were diluted with LB supplemented with 0.5M NaCl and 1 mM MgCl_2_ to 10^8^ PFU mL^-1^ and stored at 4°C until further use.

### Growth curves and infection dynamics in liquid culture

To probe growth of *E. coli* in the presence or absence of phage, cells were grown in LB until reaching an OD_600_ of 1.0, after which expression was induced with 0.1% L-arabinose. Cells were diluted with LB with 0.1% L-arabinose to an OD_600_ of 0.1 and, if indicated, infected with phage to the desired moiety of infection (MOI). Next, 100 μL of the diluted cultures were transferred to a 96-well plate and sealed with a Breathe-Easy sealing membrane (Z380059, Sigma Aldrich). Growth at 37°C, 300 rpm was tracked for 10 hours by measuring the OD_600_ at 5-minute intervals, using a Spectrostar plate reader (BMG labtech). For each measurement, two technical replicates were averaged prior to determining the average and deviations from the three independent biological replicates.

### Isolation and amplification of phage escape mutants

To isolate mutant phages, plaque assays were performed by infecting an *E. coli* strain expressing EcSduA with a 100 μL 10^8^ PFU mL^-1^ T6 phage suspension. After overnight incubation at 37°C, escapees were amplified transferring single plaques to *E. coli* cells grown in LB with 1 mM MgCl_2_ until an OD_600_ of 0.3. Growth was continued at 37°C, 220 rpm until collapse of the culture. Lysates were cleaned up as described above, after which a plaque assay was performed to verify the escape phenotype of the isolates.

### Isolation of genomic bacteriophage DNA

phage DNA was isolated using the DNeasy Blood and Tissue Kit (Qiagen) with the following adjustments. First, phages were precipitated by incubating 10 mL lysate with 8% PEG6000 for 18 hours at 4°C. Precipitated phages were centrifuged for 30 minutes at 10,000g at 4 °C, after which the supernatant was removed and the pellet was dissolved in 20 mM Tris-HCl (8.0), 250 mM NaCl to a total volume of 200 μL. Next, 20 μL of proteinase K and 200 μl AL Buffer was added and incubated for 20 minutes at 65°C. After this step, the protocol of the DNeasy Blood and Tissue Kit was followed starting from the addition of 200 μL absolute ethanol.

### Genomic sequencing and analysis of escape mutants

The genomic DNA of the escape mutants was sequenced at the Functional Genomics Center Zurich, using TruSeq Nano library preparation (Illumina) and 300 bp paired-end MiSeq sequencing (Illumina). Reads were filtered on MapQ quality using fastp^61^ with the following parameters: adapter removal, 10 bp hard trim left and right, a sliding window of 4 bp with a mapQ threshold of >20, minimum length requirement of 18 bp and overrepresentation analysis. Trimmed reads where then mapped to the T6 reference genome NC054907, using bowtie2^62^ within the Breseq analysis pipeline^63^. For the mutational analysis, the reads of the WT T6 phage were mapped to the reference genome, after which a new reference genome was assembled that harbored the pre-existing mutations in the WT T6 phage. The newly assembled reference was then used for the mutational analysis of the mutant phages with the Breseq pipeline using default settings. All silent mutations were discarded from the analysis and only mutations that were present in 100% of the reads were considered.

### Expression and purification of EcSduA

Hexahistidine-tagged SduA proteins were expressed in *E. coli* BL21-AI™ cells. Cultures were grown at 37 °C, 130 rpm until they reached an OD_600_ of 0.6, after which the cultures were incubated on ice for 1 hour. Next, protein expression was induced with 0.25 mM isopropyl-β-D-thiogalactopyranoside (IPTG), and 0.1% L-arabinose and continued for 16 h at 18 °C, 130 rpm. Cells were harvested by centrifugation and resuspended in buffer A (20 mM Tris-HCl pH 8.0, 500 mM NaCl, 5 mM imidazole, 5 mM MgCl_2_, 1 μg mL^−1^ pepstatin, 200 μg mL^−1^ AEBSF), followed by lysis in a Maximator cell homogenizer at 1,500 bar and 4 °C. The lysate was cleared by centrifugation at 10,000g for 30 min at 4 °C and applied to 15-mL equilibrated Ni-NTA beads (Qiagen). The Ni-NTA column was washed with 150 mL of buffer B (20 mM Tris-HCl pH 8.0, 500 mM NaCl, 10 mM imidazole), followed by 30 mL of buffer C (20 mM Tris-HCl pH 8.0, 150 mM NaCl, 10 mM imidazole). Proteins were eluted with five fractions of 15 mL of buffer D (20 mM Tris-HCl pH 8.0, 150 mM NaCl, 250 mM imidazole).

Protein containing elution fractions were pooled and loaded onto two equilibrated 5-mL HiTrap Heparin HP columns (Cytiva) coupled in tandem and eluted with a linear gradient of buffer E (20 mM Tris-HCl pH 8.0, 1 M NaCl). Protein containing elution fractions were pooled and dialyzed overnight against buffer F (20 mM Tris-HCl pH 8.0, 250 mM KCl, 1 mM DTT) in the presence of TEV protease. Dialyzed proteins were concentrated using 100 kDa molecular weight cut-off centrifugal filters (Merck Millipore) and further purified by size-exclusion chromatography using a Superdex 200 (16/600) column (Cytiva) equilibrated in buffer F. Purified proteins were concentrated to 10 mg mL^−1^, flash frozen in liquid nitrogen and stored at −80 °C until further use.

### Expression and purification of the N-terminal domain of EcSduA

The hexahistidine-tagged N-terminal fragment of SduA was expressed in *E. coli* BL21-AI™ cells. Cultures were grown at 37 °C, 130 rpm until they reached an OD_600_ of 0.6, after which the cultures were incubated on ice for 1 hour. Next, protein expression was induced with 0.25 mM IPTG, and 0.1% L-arabinose continued for 16 h at 18 °C, 130 rpm. Cells were harvested by centrifugation and resuspended in buffer A (20 mM Tris-HCl pH 8.0, 500 mM NaCl, 5 mM imidazole, 1 μg mL^−1^ pepstatin, 200 μg mL^−1^ AEBSF), followed by lysis in a Maximator cell homogenizer at 1,500 bar and 4 °C. The lysate was cleared by centrifugation at 10,000g for 30 min at 4 °C and applied to 15-mL equilibrated Ni-NTA beads (Qiagen). The Ni-NTA column was washed with 150 mL of buffer B (20 mM Tris-HCl pH 8.0, 500 mM NaCl, 10 mM imidazole). Proteins were eluted with five fractions of 15 mL of buffer C (20 mM Tris-HCl pH 8.0, 500 mM NaCl, 250 mM imidazole). Protein containing elution fractions were pooled and dialyzed overnight against buffer D (20 mM Tris-HCl pH 8.0, 400 mM KCl, 1 mM DTT) in the presence of TEV protease. Dialyzed proteins were concentrated using 10 kDa molecular weight cut-off centrifugal filters (Merck Millipore) and further purified by size-exclusion chromatography using a Superdex 75 (16/600) column (Cytiva) equilibrated in buffer D. Purified proteins were concentrated to 4.5 mg mL^−1^, flash frozen in liquid nitrogen and stored at −80 °C until further use.

### Crystallization of the N-terminal domain of EcSduA

Crystals of the N-terminal domain (NTD) of EcSduA were grown using the hanging drop vapor diffusion method. For crystallization, 1 μL of protein at 4.5 mg mL^−1^ was mixed with 1 μL of reservoir solution containing 100 mM HEPES pH8, 0.55 M (NH_4_)_2_SO_4_ and incubated for 39 days at 4°C. Crystals were cryoprotected by soaking in reservoir solution supplemented with 25% (v/v) glycerol and flash-frozen in liquid nitrogen. Data were collected at a temperature of 100 K at the Swiss Light Source (Paul Scherrer Institute) using the beamline PXIII and processed using autoPROC^64^. Experimental phases were obtained by the molecular replacement, using a truncated structural model of EcSduA that was predicted by the alphafold algorithm^65^. The resulting map was used for iterative atomic modelling with Coot^66^ and refinement with Phenix^67^. X-ray diffraction data and refinement statistics are provided in Supplementary Table S2.

### SEC-MALS analysis of EcSduA

For SEC-MALS purified proteins were thawed and centrifuged at 21,000g for 10 min at 4 °C, after which the supernatant was filtered with a 0.1 μm centrifugal filter (Merck Millipore). The sample was injected at a concentration 1 mg mL^-1^ onto a Superdex 200 (16/600) column (Cytiva) that was equilibrated in buffer F. After SEC, the sample passed a miniDAWN TREOS (599-TS) multiple angle light scattering detector (Wyatt Technology) and a Optilab rEX (329-rEX) refractive index detector (Wyatt Technology). The light source of the RI detector was a G1315B DAD UV detector (Agilent) and wavelength of the laser in the light scattering instrument was set at 658.9 nm. Prior to the sample a bovine serum albumin (BSA) sample was run as a calibration standard. Data collection and analysis were performed in the ASTRA 6.1 software (Wyatt Technology) with the refractive index of the solvent, the refractive index increment (∂n/∂c) and the viscosity defined as 1.331, 0.185 mL g^-1^ and 0.8945 cP, respectively.

### Sample preparation and cryo-EM data collection of apo-EcSduA

Prior to grid preparation for cryo-EM, thawed protein samples were purified by size-exclusion chromatography using a Superdex 200 (16/600) column (GE Healthcare) in equilibrated in buffer 20 mM Tris-HCl pH 8.0, 150 mM NaCl, 5 mM MgCl_2_. Peak fractions were concentrated to 2 mg mL^-1^ using 100 kDa molecular weight cut-off centrifugal filters (Merck Millipore). To each 300-mesh holey carbon grid (Au R1.2/1.3, Quantifoil Micro Tools), 2.5 μL of sample was applied and blotted for 5 s at 80% humidity and 4 °C. Grids were plunge frozen in liquid ethane, using a Vitrobot Mark IV plunger, FEI) and stored in liquid nitrogen until cryo-EM data collection. Cryo-EM data collection was performed on a FEI Titan Krios microscope equipped with a Gatan K3 direct electron detector (University of Zurich) operated at 300 kV in super-resolution counting mode. Data acquisition was performed using the EPU Automated Data Acquisition Software for Single Particle Analysis from ThermoFisher with three shots per hole at defocus range of −1.0 μm to −2.4 μm (0.2-μm steps). The final dataset comprised a total of 5,755 micrographs at a calibrated magnification of 130,000x and a super-resolution pixel size of 0.325 Å. Micrographs were exposed for 1.01s with a total dose of 66.53 e^−^ Å^−2^ over 38 subframes.

### Data processing and model building of apo-EcSduA

Cryo-EM data was processed using cryoSPARC (v3.3.2)^68^. The 5,755 micrographs were imported and motion-corrected with patch motion correction (multi) after which, the CTF values of the micrographs were estimated using patch CTF estimation (multi). Micrographs with a resolution estimate >5 Å, a defocus >2.4 μm, a total motion >92 pixels or a relative ice thickness >1.125 were discarded from the dataset (291 micrographs), yielding 5,289 micrographs for further processing steps. Next, an initial set of particles was picked with blob picker using an elliptical blob and a minimum and maximum particle diameter of 50 and 150 Å, respectively. After extraction of the particles with a box size of 384×384 pixels, particles were subjected to 2D classification to generate templates for picking (5 templates).

After template-based picking with a particle diameter of 150 Å, particles were extracted and subjected to 2D classification with a circular mask of 185 Å. Classes with defined particles were selected, resulting in a total of 708,130 particles, which were used to generate two *ab initio* models of which one was used for heterogeneous refinement with four classes. Classes were inspected visually using UCSF Chimera^69^, and the particles and volume of the best class were subjected to homogenous refinement with optimization of CTF parameters and per particle defocus enabled. The resulting particles and refined map were used as an input for non-uniform refinement with optimization of CTF parameters and C2 symmetry enabled. The final map was sharpened with a B-factor of -75. The local resolution was estimated based on the resulting map using the local resolution function of cryoSPARC and plotted on the map using UCSF Chimera^69^.

The structural model of EcSduA was built *de novo* in Coot (V0.9.2)^66^ and was refined over multiple rounds using Phenix^67,70^. Real-space refinement was performed with the global minimization, atomic displacement parameter (ADP) refinement and secondary structure restrains enabled. The quality of the atomic model, including protein geometry, Ramachandran plots, clash analysis and model cross-validation, was assessed with MolProbity and the validation tools in Phenix^67,70–72^. The refinement statistics of the final model are listed in Supplementary Table S1. Figures of maps, models and the calculations of map contour levels were generated using ChimeraX^69^.

### Sample preparation and cryo-EM data collection on the EcSduA DNA-bound complex

Prior grid preparation for cryo-EM, thawed protein samples were incubated on ice for 20 minutes with a 40 bp dsDNA substrate at a 1.0:1.3 molar ratio (protein: DNA). Next, the sample was purified by size-exclusion chromatography using a Superdex 200 (16/600) column (GE Healthcare) in equilibrated in buffer 20 mM Tris-HCl pH 8.0, 150 mM NaCl. Peak fractions were concentrated to 2.2 mg mL^-1^ using 100 kDa molecular weight cut-off centrifugal filters (Merck Millipore). To each 300-mesh holey carbon grid (Au R1.2/1.3, Quantifoil Micro Tools), 2.5 μL of sample was applied and blotted for 5 s at 80% humidity and 4 °C. Grids were plunge frozen in liquid ethane, using a Vitrobot Mark IV plunger, FEI) and stored in liquid nitrogen until cryo-EM data collection. Cryo-EM data collection was performed on a FEI Titan Krios microscope equipped with a Gatan K3 direct electron detector (University of Zurich) operated at 300 kV in super-resolution counting mode. Data acquisition was performed using the EPU Automated Data Acquisition Software for Single Particle Analysis from ThermoFisher with three shots per hole at defocus range of −1.0 μm to −2.4 μm (0.2-μm steps). The final dataset comprised a total of 5,755 micrographs at a calibrated magnification of 130,000x and a super-resolution pixel size of 0.325 Å. Micrographs were exposed for 1.01s with a total dose of 60.01 e^−^ Å^−2^ over 38 subframes.

### Data processing and model building of the EcSduA DNA-bound complex

Cryo-EM data was processed using cryoSPARC (v4.1.1)^68^. The 9,799 micrographs were imported and motion-corrected with patch motion correction (multi) after which, the CTF values of the micrographs were estimated using patch CTF estimation (multi). Micrographs with a resolution estimate >5 Å, a defocus >2.4 μm or a total motion >50 pixels were discarded from the dataset (604 micrographs), yielding 9,195 micrographs for further processing steps. Next, an initial set of particles was picked with blob picker using an elliptical blob and a minimum and maximum particle diameter of 50 and 150 Å, respectively. After extraction of the particles with a box size of 384×384 pixels, particles were subjected to 2D classification to generate templates for picking (5 templates).

After template-based picking with a particle diameter of 150 Å, particles were extracted and subjected to 2D classification with a circular mask of 185 Å. Classes with defined particles were selected, resulting in a total of 1,064,688 particles, which were used to generate two *ab initio* models of which one was used for heterogeneous refinement with four classes. Classes were inspected visually using UCSF Chimera^69^, and the particles and volume of the best class were subjected to 3D variability analysis with 4 clusters. The resulting volume series were inspected visually using UCSF Chimera^69^ and volume with the best features was used for heterogeneous refinement with three classes. The resulting particles and refined map were used as an input for non-uniform refinement with optimization of CTF parameters and optimization of per-particle defocus enabled. The final map was sharpened with a B-factor of -80. The local resolution was estimated based on the resulting map using the local resolution function of cryoSPARC and plotted on the map using UCSF Chimera^69^.

The structural model of EcSduA bound to DNA was built in Coot (V0.9.2)^66^ and was refined over multiple rounds using Phenix^67,70^. Real-space refinement was performed with the global minimization, atomic displacement parameter (ADP) refinement and secondary structure restrains enabled. The quality of the atomic model, including protein geometry, Ramachandran plots, clash analysis and model cross-validation, was assessed with MolProbity and the validation tools in Phenix^67,70–72^. The refinement statistics of the final model are listed in Supplementary Table 1. Figures of maps, models and the calculations of map contour levels were generated using UCSF Chimera^69^.

### *In vitro* nuclease activity assays with fluorescently labelled oligo nucleotides

Prior to the cleavage assays, synthetic dsDNA targets (Integrated DNA technologies) were annealed in a buffer containing 10 mM Tris-HCl, 50 mM NaCl, using a thermocycler (Biorad) and stored at -20°C until further use. For the cleavage assays, 40 nM of DNA or RNA target was mixed with 400 nM of the EcSduA complex in a buffer containing 20 mM Tris-HCl, 100 mM NaCl and 5 mM MgCl_2_ and incubated for 1 hour at 37°C, unless stated otherwise. After the incubation, the reaction was quenched by the addition of 1 μL proteinase K and incubated for 10 minutes at 60°C. Next, samples were mixed with loading dye (97% formamide, 25 mM EDTA and 0.15% Orange G), incubated for 10 minutes at 95°C and loaded onto a 15% or 10% denaturing PAGE gel containing 8M urea. Gels were run for 2.5 hours at 350 V, followed by imaging with the Typhoon trio (GE healthcare). All sequences of oligonucleotides used in this study are provided in Supplementary Table 3.

### *In vitro* cleavage assays with genomic phage DNA

For the cleavage assays with genomic DNA, 100 pM of genomic DNA was mixed with 10 nM of the EcSduA complex in a buffer containing 20 mM Tris-HCl, 100 mM NaCl and 5 mM MgCl_2_ and incubated for the indicated times at 37°C. After the incubation, the reaction was quenched by the addition of 1 μL proteinase K. Next, samples were mixed with 6x DNA loading dye (10 mM Tris-HCl (pH 7.6), 60 mM EDTA, 60% Glycerol, 0.03% Bromophenol blue and 0.03% Xylene cyanol FF) and loaded onto a 1% agarose gel. Gels were run for 40 minutes at 100 V, followed by imaging with an ChemiDoc Imaging System (Biorad).

### Fluorescence polarization binding assays

For fluorescence polarization binding assays, 20 nM of DNA or RNA substrate was mixed with various concentrations of the EcSduA complex in a binding buffer containing 20 mM Tris-HCl, 250 mM NaCl, 0.05% Tween20 and incubated for 20 minutes at room-temperature. After the incubation, samples were transferred to a 384 well plate and fluorescence polarization was measured using a Pherastar plate reader (BMG labtech). For each measurement, the experiment was repeated three independent times to determine the average and deviations.

### Plasmid transformation assays

Prior to the transformation assays, Mach1 *E. coli* cells carrying an empty pET plasmid or an EcSduA variant were made chemically competent. To make cells chemically competent, cultures were grown in 200 mL of LB supplemented with 50 μg mL^-1^ kanamycin and 10 μM IPTG Cultures were grown at 37 °C, 130 rpm until they reached an OD_600_ of 0.4, after which the cultures were incubated on ice for 30 minutes. Cells were harvested by centrifugation for 10 minutes at 4000g. The supernatant was discarded, and the pellet was dissolved with 25 mL of ice cold 100 mM MgCl_2_, followed by centrifugation for 10 minutes at 4000g. The pellet was dissolved in 25 mL of ice cold 100 mM CaCl_2_ and incubated for 20 minutes on ice. Next, cells were harvested by centrifugation for 10 minutes at 4000g and the pellet was dissolved with 8 mL of ice cold 85 mM MgCl_2_ supplemented with 15% glycerol, after which 140 μL of DMSO was added to the cells. The cells were incubated on ice for 15 minutes, aliquoted, frozen in liquid nitrogen and stored at -80°C until further use.

DNA templates for the transformation assays were prepared by PCR amplification from the pUC19 plasmid (NEB), after which the PCR fragments were Dpn1-treated, and gel purified using the Qiagen NucleoSpin Gel cleanup kit (Qiagen). For transformation assays with DNA fragments, the backbone was mixed with the insert at a 1:5 ratio, after which 10 ng DNA was incubated for 30 minutes on ice with 50 μL of chemically competent cells. After incubation, cells were incubated at 42°C for 45 seconds, followed by 3 minutes incubation on ice. Subsequently, 500 μL of LB was added to the cells and incubated for 1.5 hours at 37°C, 600 rpm. Next, 100 μL of cells were plated onto LB agar plates containing 50 μg mL^-1^ kanamycin, 100 μg mL^-1^ ampicillin and 100 μM IPTG and incubated at 37 °C for 18 hours, after which colonies were counted. Colony forming units were normalized against the average counts of the vector only control.

