## Supplemental information for "DNA end sensing and cleavage by the Shedu anti-phage defense system"

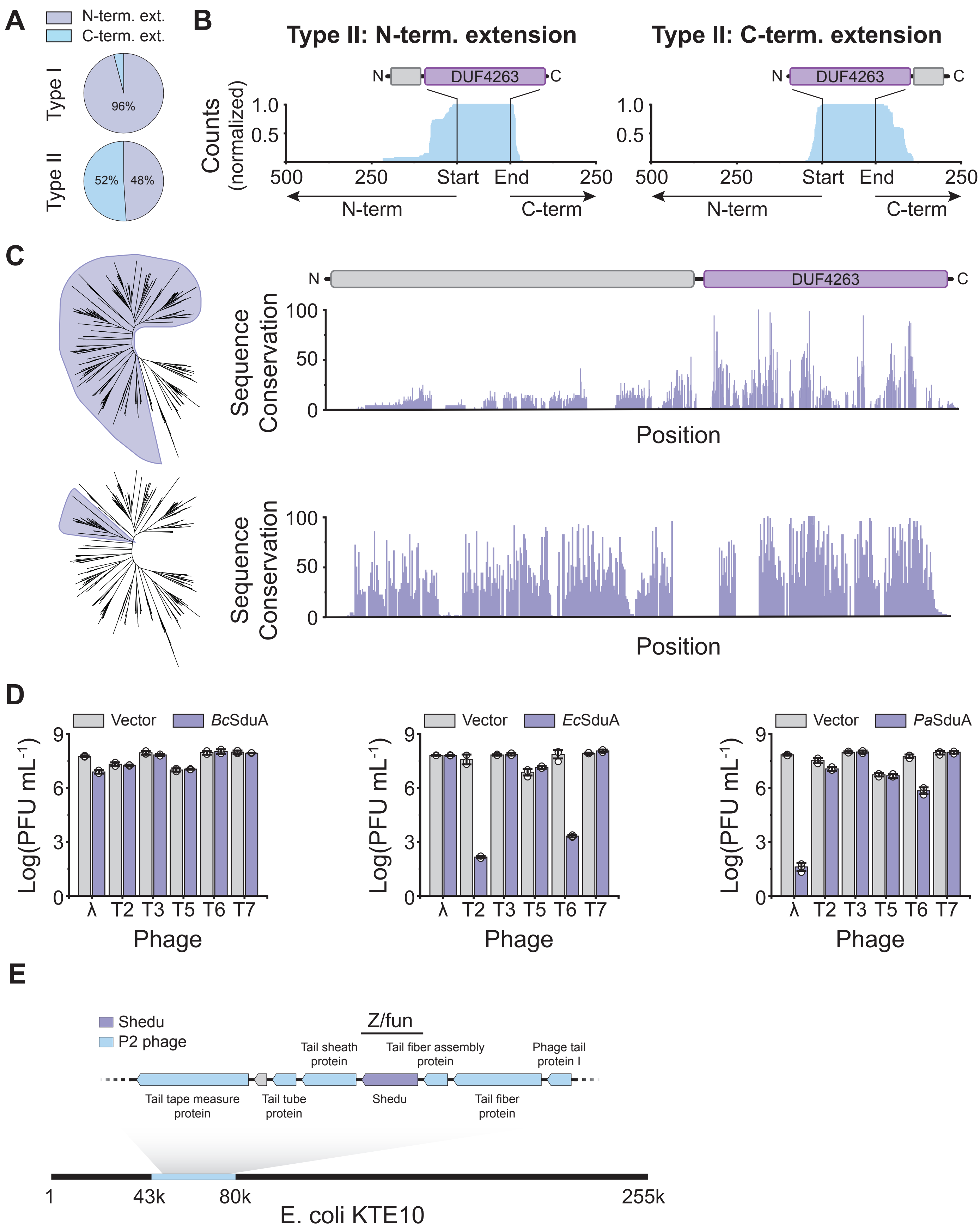


**Figure S1: Bioinformatic analysis of Shedu anti-phage systems, related to Figure 1.**

**(A)** Distribution of SduA proteins containing N- or C-terminal extensions of the DUF4263 domain, within the type I and type II systems. **(B)** Architecture of type II SduA orthologs with either N- or C-terminal extensions. **(C)** Sequence conservation within all type I systems (top) and a specific clade within the type I systems (bottom). **(D)** Plaque assays of *E. coli* strains expressing SduA orthologs using a panel of phages. Data represents mean PFU mL$^{-1}$ ± SEM of three independent replicates (n=3). **(E)** Locus organization of *E. coli* KTE10 strain that harbors the EcSduA-containing system within the Z/Fun locus of a P2 prophage.

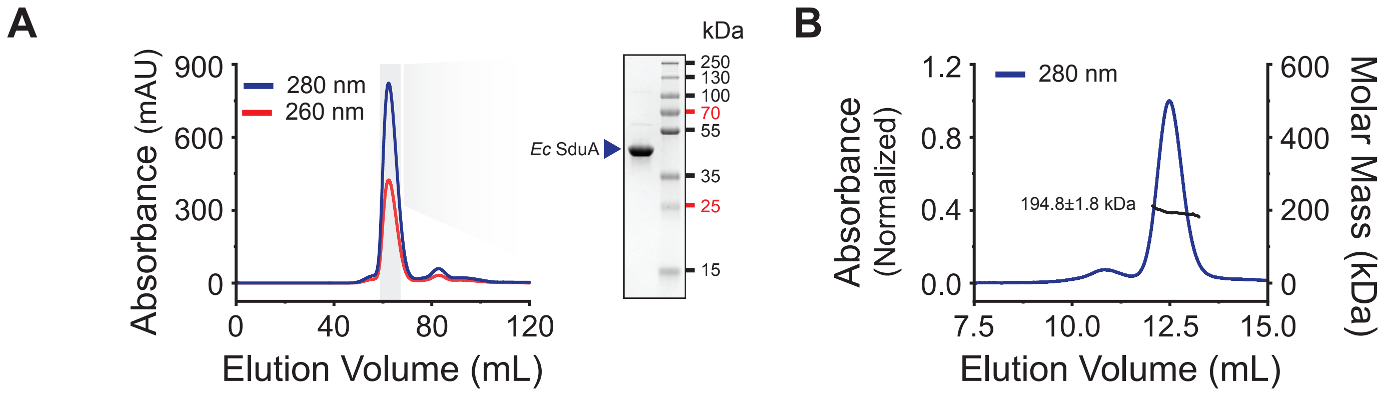


**Figure S2: Purification and size determination for the EcSduA complex, related to Figure 2.**

**(A)** Analysis of purified recombinantly-epressed EcSduA by size-exclusion chromatography (left) and SDS-PAGE (right). **(B)** Size-exclusion chromatography coupled to multi-angle static light scattering (SEC-MALS) analysis of the EcSduA complex. The calculated molecular mass indicates a tetrameric assembly.

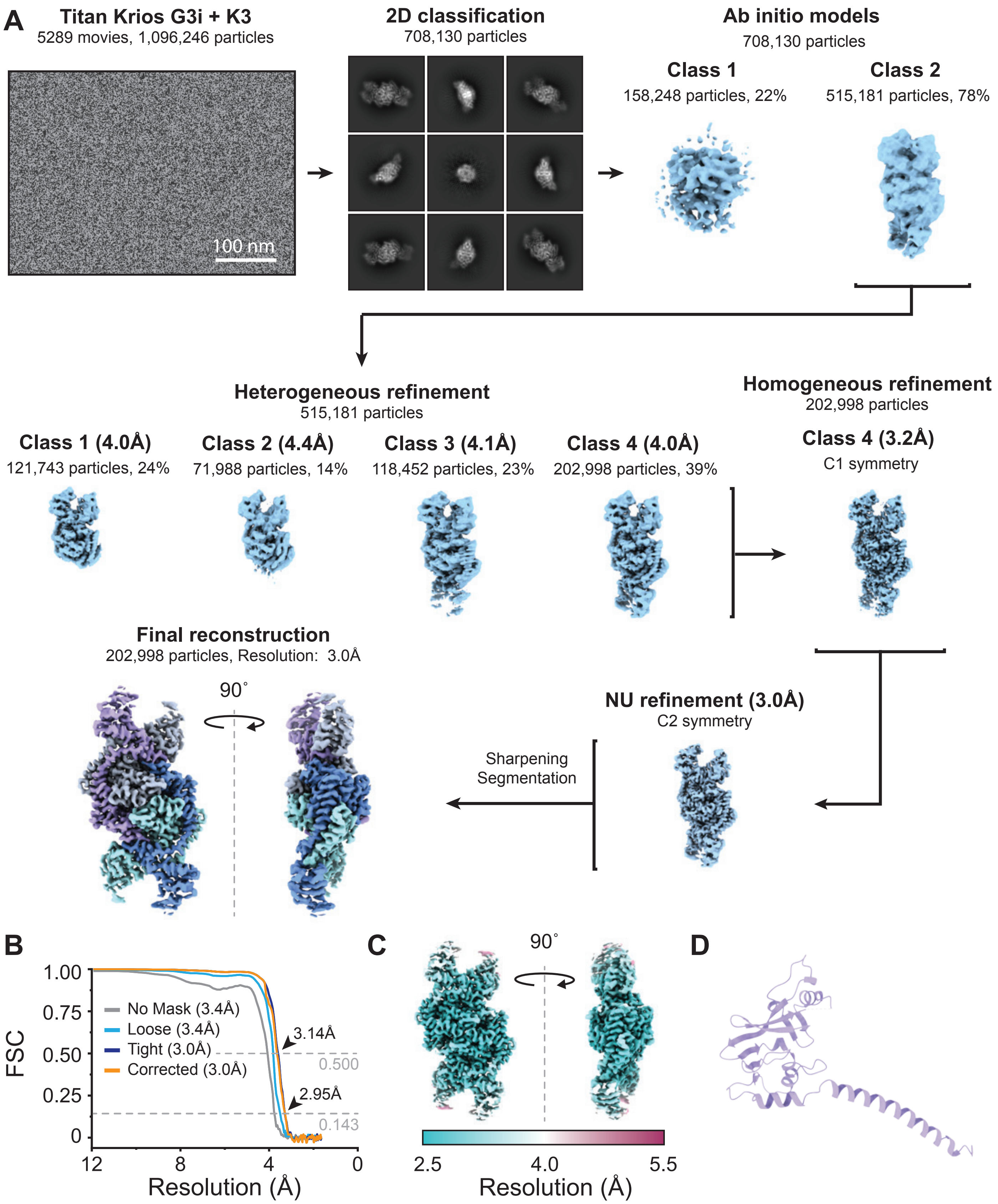


**Figure S3: Cryo-EM processing workflow for the EcSduA complex and crystal structure of the N-terminal domain of EcSduA, related to Figure 2.**

**(A)** Cryo-EM processing workflow for the EcSduA-dsDNA complex. **(B)** Fourier Shell Correlation (FCS) determined from two independently refined half-maps. The gold standard cut-off (FCS=0.143) is marked with an arrow. **(C)** Local resolution estimation on the final cryo-EM density map of the EcSduA complex. **(D)** Crystal structure of N-terminal domain of EcSduA, determined at a resolution of 2.5 Å.

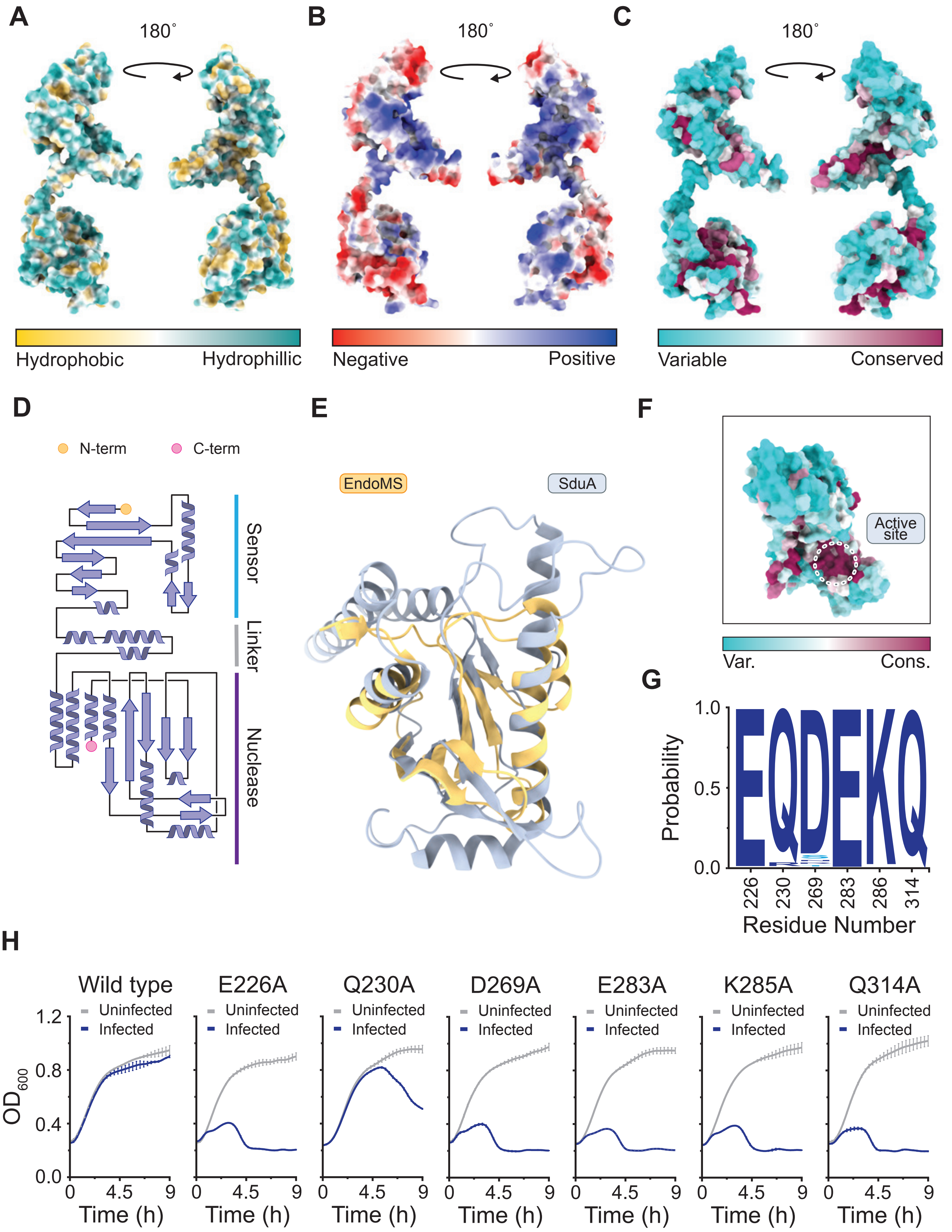


**Figure S4: Molecular details of the EcSduA monomer, related to Figure 2 and 3.**

**(A)** Surface view of an EcSduA monomer colored by hydrophobicity. **(B)** Surface view of EcSduA, shown in the same orientation as in (A), colored by electrostatic potential. **(C)** Surface view of EcSduA, shown in the same orientation as in (A), colored by sequence conservation. **(D)** Topology diagram of the secondary structure of EcSduA. **(E)** Structural superposition of the C-terminal domain of EcSduA (light grey) with the PD/ExK nuclease domain of EndoMS (orange,

PDB: 5GKH)[55]. **(F)** Top view of an EcSduA monomer colored by sequence conservation. The nuclease active site is indicated by a dotted circle. **(G)** Sequence conservation analysis of active site residues in Shedu. Figure was generated using WebLogo[56]. **(H)** Growth curves of *E. coli* strains expressing WT EcSduA or active site mutants. Infected curves represent cultures that were infected with the T6 phage at an MOI of 5. Data points represent the mean ± SEM of three biological replicates (n=3).

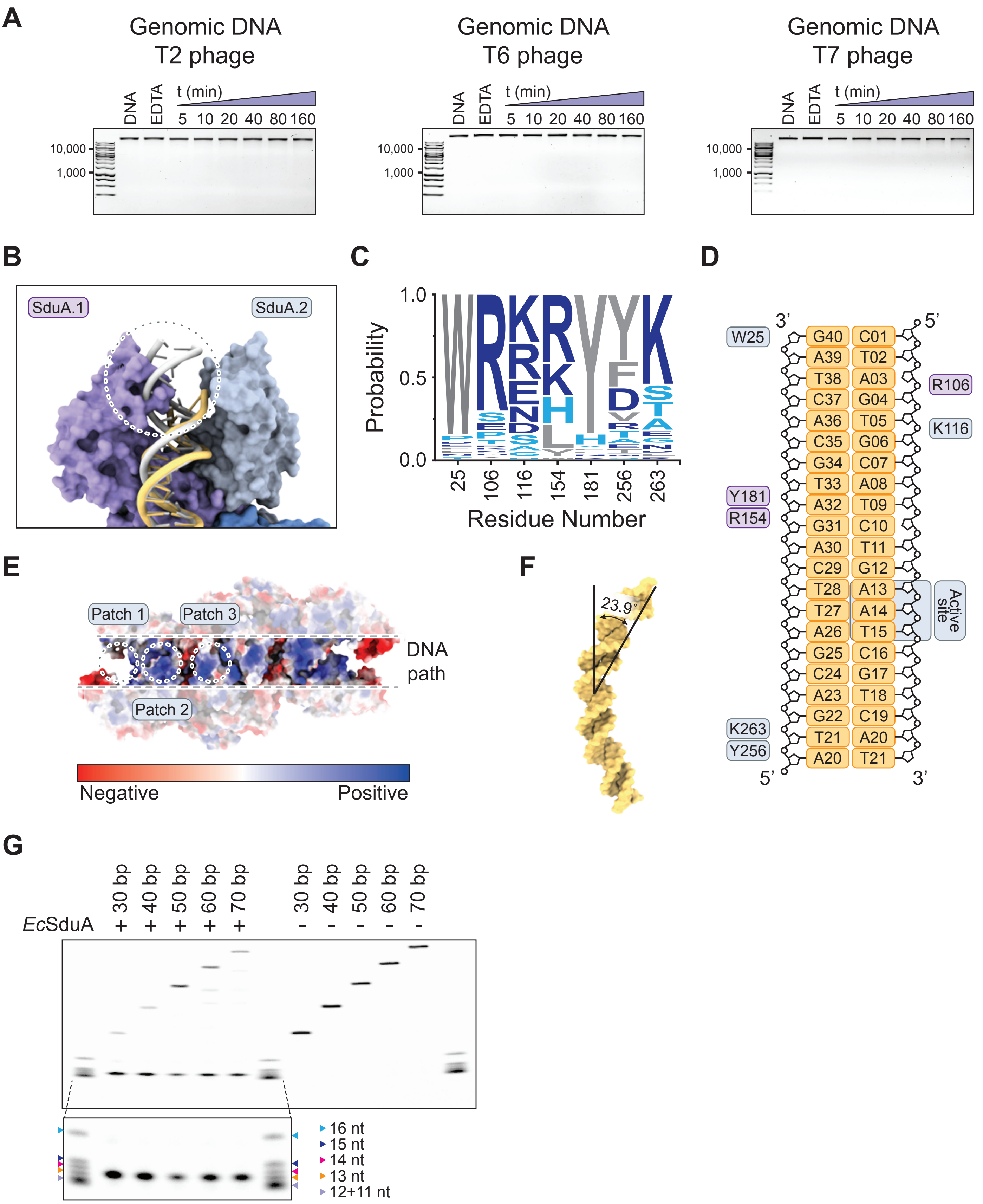


**Figure S5: Molecular details of the DNA end recognition by EcSduA, related to Figure 4 and 5.**

**(A)** Analysis of nuclease activity of EcSduA on genomic DNA isolated from T2, T6 and T7 phages. T2 and T6 extensively modify their DNA by glucosylation[30,31], whereas the DNA of T7 is unmodified. **(B)** Structural superposition of the dsDNA substrate bound by the EcSduA complex (light orange) and an ideal B-form dsDNA substrate (light gray). Steric clashes are indicated by a dotted circle. **(C)** Sequence conservation analysis of residues in the N-terminal clamp of Shedu. Figure was generated using WebLogo[56]. **(D)** Schematic showing the residues that interact with the dsDNA substrate. **(E)** Surface view of the EcSduA complex colored by electrostatic potential. Interaction interfaces, called Patch 1 to 3, are indicated by dotted circles. **(F)** Kinked geometry of dsDNA bound to EcSduA. **(G)** Re-analysis of samples in Figure 5C displaying of nicking activity of WT EcSduA on fluorescently labelled dsDNA oligonucleotides of increasing lengths. Nicking products were resolved by denaturing PAGE alongside fluorescently labelled oligonucleotides of a defined length.

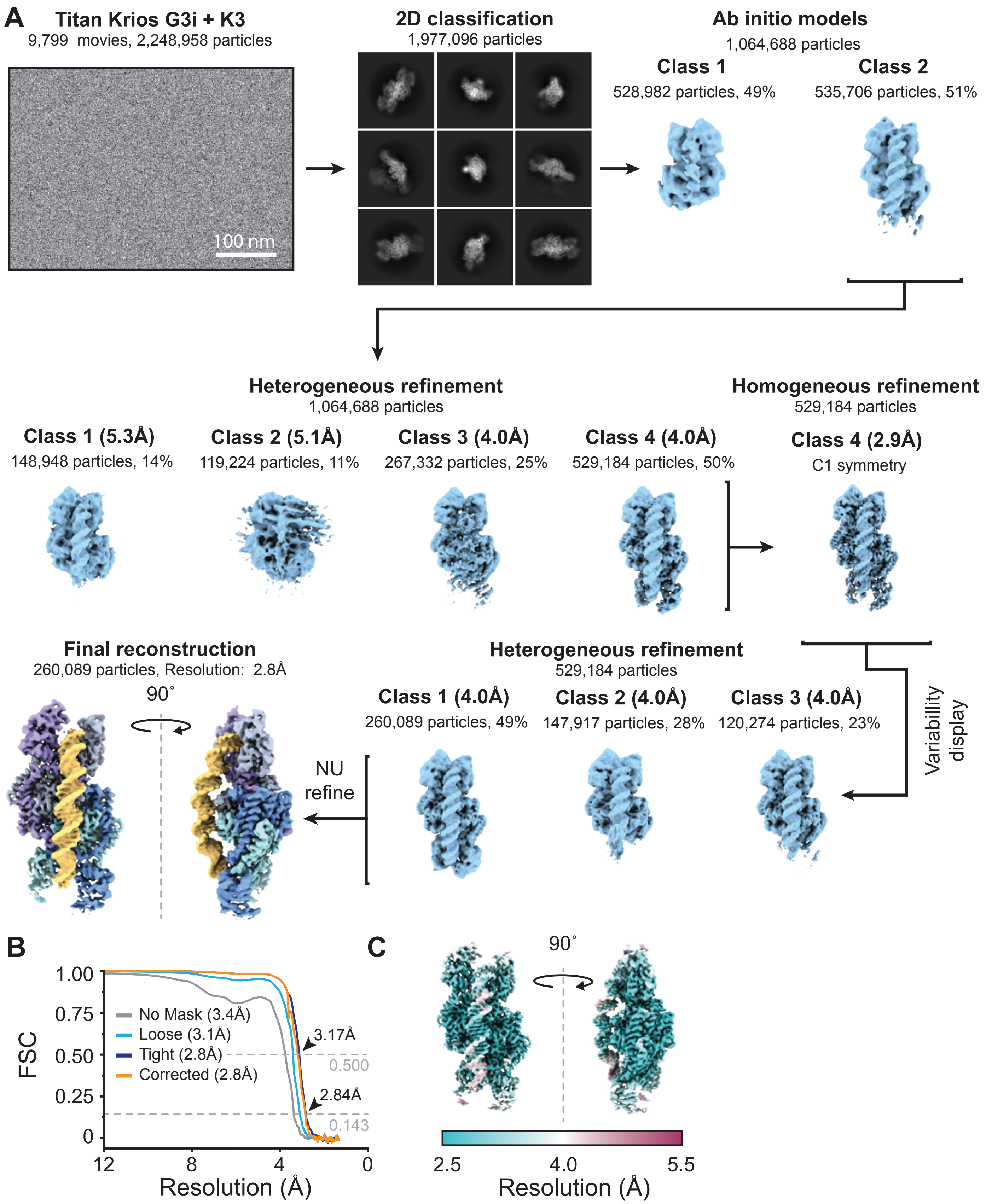


**Cyo-EM processing workflow for EcSduA in complex with a dsDNA substrate, related to Figures 4 and 5.**

**(A)** Cryo-EM processing workflow for the EcSduA-dsDNA substrate complex. **(B)** Fourier Shell Correlation (FCS) determined from two independently refined half-maps. The gold standard cut-off (FCS=0.143) is marked with an arrow. **(C)** Local resolution estimation on the final cryo-EM density map of the EcSduA complex.

## A

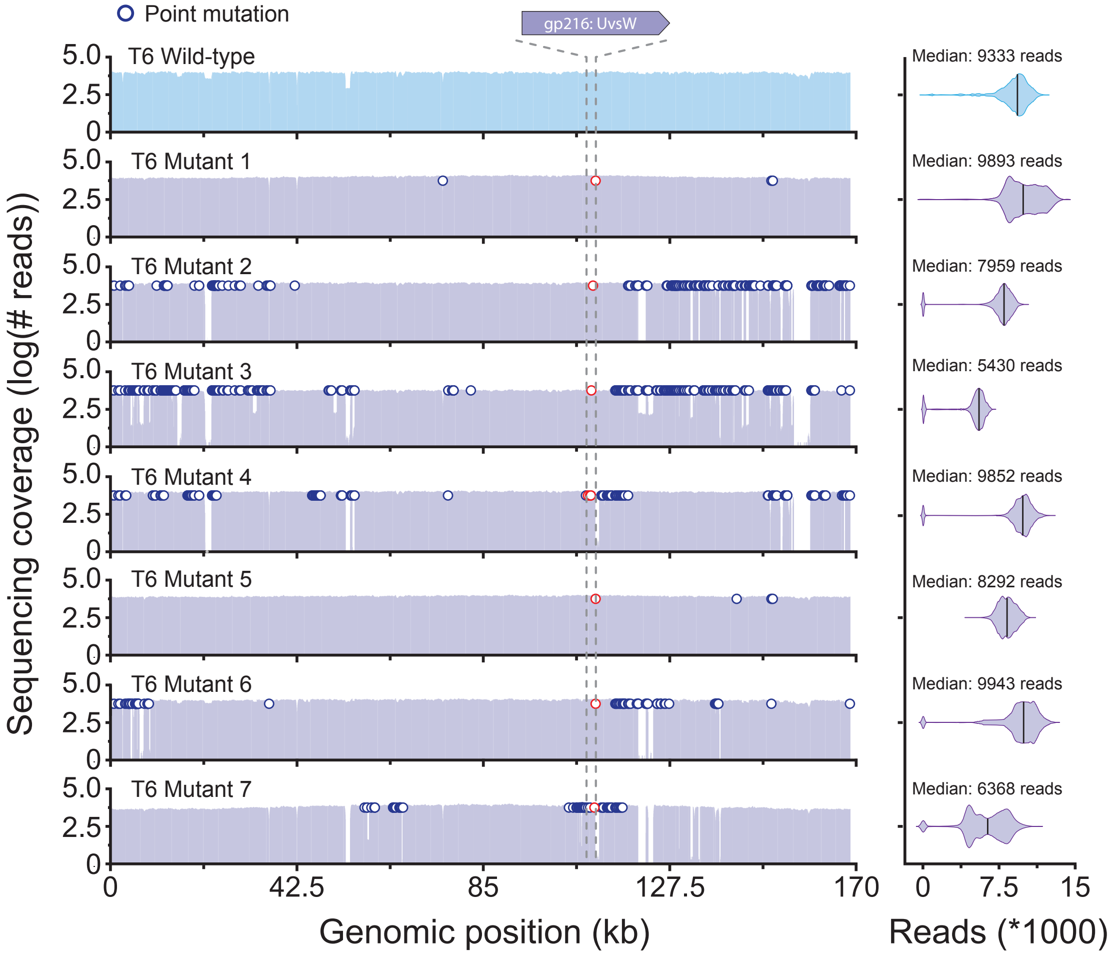


## B

| Phage | Position | Mutation | Gene | Effect |
|---|---|---|---|---|
| Mutant 1 | 110,608 | (A)deletion | gp216: UvsW | Frame shift (1410/1509 nt) |
| Mutant 2 | 110,009 | C → T | gp216: UvsW | Pre-mature stop (Q271*) |
| Mutant 3 | 109,644 | C → A | gp216: UvsW | Mutation (A149E) |
| Mutant 4 | 109,329 | G → A | gp216: UvsW | Mutation (C44Y) |
| Mutant 4 | 109,494 | G → A | gp216: UvsW | Pre-mature stop (W99*) |
| Mutant 4 | 109,496 | C → A | gp216: UvsW | Mutation (L100I) |
| Mutant 4 | 109,504 | A → T | gp216: UvsW | Mutation (K102N) |
| Mutant 5 | 110,608 | (A)insertion | gp216: UvsW | Frame shift (1410/1509 nt) |
| Mutant 6 | 110,608 | (A)deletion | gp216: UvsW | Frame shift (1410/1509 nt) |
| Mutant 7 | 109,329 | G → A | gp216: UvsW | Mutation (C44Y) |
| Mutant 7 | 110,608 | G → T | gp216: UvsW | Mutation (G379V) |

**Figure S6: Whole genome sequencing of EcSduA escaper phages, related to Figure 5.**

**(A)** Genome coverage of T6 WT and escaper phages (left). The positions of uncorrelated point mutations are indicated by purple circles, whereas correlated point mutations in *uvsW* are indicated by red circles. Violin plots showing the read depth, median is indicated by the solid black line. **(B)** Overview of *uvsW* mutations identified in the escaper phages.

**Table S1: Cryo-EM data collection, refinement, and validation statistics for the *Escherichia coli* SduA complexes**

| | *Escherichia coli* SduA Complex. (EMDB: EMD-17844) (PDB 8PS4) | *Escherichia coli* SduA complex bound to a dsDNA substrate. (EMDB: EMD-17845) (PDB 8PS5) |
|---|---|---|
| **Data collection and processing** | | |
| Magnification | 130,000 x | 130,000 x |
| Voltage (kV) | 300 | 300 |
| Electron exposure (e–/Å$^2$) | 66.53 | 60.01 |
| Defocus range (µm) | -1.0 to -2.4 (-0.2 steps) | -1.0 to -2.4 (-0.2 steps) |
| Pixel size (Å) | 0.65 | 0.65 |
| Symmetry imposed | C2 | C1 |
| Initial particle images (no.) | 1,096,246 | 2,248,958 |
| Final particle images (no.) | 202,998 | 260,089 |
| Map resolution (Å) | 2.95 | 2.84 |
| FSC threshold | 0.143 | 0.143 |
| Map resolution range (Å) | 2.5 – 5.5 | 2.5 – 5.5 |
| **Refinement** | | |
| Model resolution (Å) | 2.9 | 2.7 |
| FSC threshold | 0.143 | 0.143 |
| Model resolution range (Å) | 2.6-3.2 | 2.5-3.1 |
| Map sharpening *B* factor (Å$^2$) | -75 | -80 |
| Model composition | | |
| Non-hydrogen atoms | 12310 | 14866 |
| Protein residues | 1616 | 1624 |
| Nucleotides | 0 | 80 |
| Ligands | MG 2 | - |
| *B* factors (Å$^2$) | | |
| Protein | 30.00/238.95/114.86 | 18.43/260.46/95.66 |
| Nucleotides | - | 26.48/294.67/145.49 |
| Ligand | 30.00/30.00/30.00 | - |
| R.m.s. deviations | | |
| Bond lengths (Å) | 0.012 | 0.006 |
| Bond angles (°) | 1.464 | 0.938 |
| Validation | | |
| MolProbity score | 1.64 | 1.66 |
| Clashscore | 7.52 | 6.98 |
| Poor rotamers (%) | 1.97 | 1.49 |
| Ramachandran plot | | |
| Favored (%) | 98.25 | 97.21 |
| Allowed (%) | 1.75 | 2.79 |
| Disallowed (%) | 0.00 | 0.00 |

**Table S2: X-ray crystallographic data collection and refinement statistics for the N-terminal fragment of *Escherichia coli* SduA**

| | N-terminal fragment of *Escherichia coli* SduA (PDB: 8PS6) |
|---|---|
| **Data collection** | |
| Space group | R 3 2 :H |
| Cell dimensions | |
| *a*, *b*, *c* (Å) | 225.543 225.543 32.029 |
| α, β, γ (°) | 90 90 120 |
| Resolution (Å) | 37.59 - 2.52 (2.56 - 2.52) |
| $R_{sym}$ or $R_{merge}$ | 0.1713 (1.4) |
| $I$ / σ$I$ | 13.99 (2.45) |
| Completeness (%) | 95.74 (100.00) |
| Redundancy | 20.3 (21.3) |
| **Refinement** | |
| Resolution (Å) | 37.59 - 2.52 |
| No. reflections | 10117 (1048) |
| $R_{work}$ / $R_{free}$ | 22.29 / 0.2434 |
| No. atoms | |
| Protein | 1485 |
| Ligand/ion | 10 |
| Water | 33 |
| *B*-factors | |
| Protein | 59.65 |
| Ligand/ion | 107.32 |
| Water | 52.65 |
| R.m.s. deviations | |
| Bond lengths (Å) | 0.009 |
| Bond angles (°) | 1.20 |

**Table S3: Oligonucleotides used in this study.**

| Identifier | Sequence (5' --> 3') | Description | Figure |
|---|---|---|---|
| oLL112 | AAAAAAAAAAGTGACTATTCAAGCATACAGGCTGATTAAC | ssDNA substrate, 5' Atto647NN labeled | 3A, B |
| oAI010 | UAAAUCAGAUUGGAAUCACUGCUAUGCAGCUUAUUC | ssRNA substrate, 5' Atto532 labeled | 3A, B |
| oLL321 | GATACCAGCTCTATGTTGACCTCTCAGATCGGCACTACTT | 40 bp dsDNA substrate, 5' Cy5 labeled | 3A, B |
| oLL322 | AAGTAGTGCCGATCTGAGAGGTCAACATAGAGCTGGTATC | 40 bp dsDNA substrate, unlabeled | 3A, B, E |
| oLL314 | CTAGTGCATCTGAATCGTCATGACGATTCAGATGCACTAG | 40 bp palindromic substrate for cryoEM | 4A |
| oLL323 | GATACCAGCTCTATGTTGACCTCTCAGATC | 30 bp dsDNA substrate, 5' Cy5 labeled | 5C |
| oLL324 | GATCTGAGAGGTCAACATAGAGCTGGTATC | 30 bp dsDNA substrate, unlabeled | 5C |
| oLL319 | GATACCAGCTCTATGTTGACCTCTCAGATCGGCACTACTTGAGTCCAATC | 50 bp dsDNA substrate, 5' Cy5 labeled | 5C |
| oLL320 | GATTGGACTCAAGTAGTGCCGATCTGAGAGGTCAACATAGAGCTGGTATC | 50 bp dsDNA substrate, unlabeled | 5C |
| oLL317 | GATACCAGCTCTATGTTGACCTCTCAGATCGGCACTACTTGAGTCCAATCTATTTGAAGC | 60 bp dsDNA substrate, 5' Cy5 labeled | 5B, C |
| oLL318 | GCTTCAAATAGATTGGACTCAAGTAGTGCCGATCTGAGAGGTCAACATAGAGCTGGTATC | 60 bp dsDNA substrate, unlabeled | 5C |
| oLL336 | GCTTCAAATAGATTGGACTCAAGTAGTGCCGATCTGAGAGGTCAACATAGAGCTGGTATC | 60 bp dsDNA substrate, 3' Cy3 labeled | 5B |
| oLL316 | GATACCAGCTCTATGTTGACCTCTCAGATCGGCACTACTTGAGTCCAATCTATTTGAAGCTACCATGGAC | 70 bp dsDNA substrate, 5' Cy5 labeled | 5C |
| oLL317 | GTCCATGGTAGCTTCAAATAGATTGGACTCAAGTAGTGCCGATCTGAGAGGTCAACATAGAGCTGGTATC | 70 bp dsDNA substrate, unlabeled | 5C |
| oLL341 | GACAAGCTGTGACCGTCTCC | pUC19 Back Bone primer Forward | 5E |
| oLL342 | CTTGGCGTAATCATGGTCATAGCT | pUC19 Back Bone primer Reverse | 5E |
| oLL343 | ATGACCATGATTACGCCAAGCTTG | pUC19 Insert Forward | 5E |
| oLL344 | GGAGACGGTCACAGCTTGTC | pUC19 Insert Reverse | 5E |
